# Mutual Inhibition Model of pattern formation: The role of Wnt-Dickkopf interactions in driving *Hydra* body axis formation

**DOI:** 10.1101/2021.09.13.460125

**Authors:** Moritz Mercker, Alexey Kazarnikov, Anja Tursch, Thomas Richter, Suat Özbek, Thomas Holstein, Anna Marciniak-Czochra

## Abstract

The antagonistic interplay between canonical Wnt signalling and Dickkopf (Dkk) proteins is fundamental to tissue organisation, including stem cell differentiation and body-axis formation. Disruptions in this interaction are linked to various human diseases, yet the mechanisms enabling robust body-axis formation through *β*-catenin/Wnt–Dkk interactions remain unclear. A key model system for Wnt-driven pattern formation is the pre-bilaterian organism *Hydra*, where two ancestral Dkk proteins interact with Wnt signalling to self-organise the body axis. While *Hydra* patterning has been extensively studied using the activator–inhibitor framework, a model integrating experimentally identified molecules has been lacking. Here, we introduce a mathematical model that incorporates both Dkks and their experimentally observed interactions with Wnt signalling. Numerical and analytical studies show that this network alone is sufficient to drive *de novo* body-axis formation across a broad parameter range. Our mutual inhibition model provides a biologically grounded realization of the general local-activation/long-range-inhibition (LALI) principle of *de novo* pattern formation, offering a mechanistic explanation for the observed *Dkk* and *Wnt* expression patterns under various conditions. Unlike previous models, it is directly grounded in experimental data, links injury response to pattern formation, and remains robust against perturbations.

**Author Summary:** How organisms form and regenerate complex body structures is a fundamental question in biology. In the freshwater animal *Hydra*, which can regenerate its entire body from a small tissue fragment, a molecular signalling system involving Wnt proteins and their inhibitors, the Dickkopf (Dkk) family, plays a central role in organising the body axis. While these molecules are known to interact, their exact roles and how they collectively shape large-scale patterns have remained unclear—especially since their activity does not fully align with established pattern formation models. In this study, we present a new mathematical model that captures the observed interactions between Wnt and two Dkk molecules in *Hydra*. We show that a mechanism based on mutual inhibition—rather than the traditional interplay between activator and inhibitor molecules—can explain the emergence of a stable body axis and the results of various perturbation experiments. Our work offers new insights into the design principles of biological pattern formation and emphasizes the importance of exploring alternative mechanisms beyond classical theories.

## Introduction

Cnidarians, with their simple body plans and remarkable regenerative abilities, offer a powerful model system for studying fundamental and broadly applicable principles of development and pattern formation [1–3]. Among them, *Hydra* has served as a classic organism in developmental biology for nearly 300 years, owing to its continuous morphogenetic activity, capacity for whole-body regeneration, and amenability to experimental manipulation [4–7]. In *Hydra*, body axis formation is a striking example of a self-organising process: even when dissociated into individual cells, aggregates can regenerate into functional polyps [8, 9]. This regeneration showcases *de novo* pattern formation, where cells determine their fate based on positional cues rather than retaining memory of their axial origin [10, 11].

To explain such processes, various mathematical models have been proposed. Many adopt a top-down, abstract approach to infer the interactions that could underlie observed patterns. A foundational concept comes from Alan Turing’s theory of reaction-diffusion systems, where interacting morphogens produce patterns through diffusion-driven instabilities [12–14]. Building on this, Gierer and Meinhardt developed the activator-inhibitor model for *Hydra*, postulating that local self-enhancement of an activator, coupled with long-range inhibition, could account for head, foot, and tentacle patterning [15–17]. A later refinement introduced the concept of a slowly changing source density (SD), representing a memory of the body axis that stabilises and aligns spatial domains [18, 19].

From a molecular perspective, canonical Wnt/*β*-Catenin signalling governs posterior identity in many organisms, while inhibitors such as Dickkopf (Dkk) proteins define anterior fates by antagonising Wnt activity [20–24]. In vertebrates, Dkk proteins play essential roles in axial and head development and have been linked to various human diseases, including cancer and neurodegeneration [21, 25]. In *Hydra*, a similar antagonism between Wnt/*β*-Catenin signalling and *HyDkk* genes appears to underlie axis and head formation during regeneration [4, 5, 26–30]. Nuclear *β*-Catenin and expression of genes like *HyWnt3* mark the oral pole, while *HyDkk1/2/4-A* and *HyDkk1/2/4-C* are expressed in the body column and suppress Wnt signalling downstream [29, 30]. These genes are considered evolutionary precursors of vertebrate Dkk homologs [29].

However, while the activator-inhibitor model provides a compelling theoretical framework for *de novo* pattern formation, its alignment with known molecular pathways in *Hydra* remains problematic. Although canonical *HyWnt* signalling is a plausible candidate for the activator, a corresponding diffusible inhibitor that fits the model’s assumptions has yet to be identified. Moreover, regeneration in *Hydra* often requires strong wound signals to initiate pattern formation, rather than emerging spontaneously from instability in a homogeneous field [19,29,31]. This challenges the model’s central premise of self-organisation without external cues.

Expression patterns of *HyDkk* genes in *Hydra* deviate from the activator-inhibitor model predictions. *HyDkk1/2/4-A* is uniformly expressed, and *HyDkk1/2/4-C* decreases towards the aboral end—neither is expressed at the oral pole, contradicting the expected inhibitor gradient [29, 30]. Additionally, both genes are downregulated upon GSK3 inhibition, complicating their role in Wnt regulation and pattern formation [29, 30]. These discrepancies led to a reconsideration of their function, and the molecules were excluded as candidates for the self-organised pattern formation framework in *Hydra* [17, 32]. Consequently, research shifted towards identifying a missing inhibitor to fit the activator-inhibitor model, such as the transcription factor *Sp5*. Yet *Sp5* lacks diffusibility and fails to match predicted spatial expression profiles [33].

Other candidates, including thrombospondin (*TSP*) and the secreted protease *HAS-7*, have been proposed as Wnt antagonists [34, 35]. However, their expression patterns and functional effects do not conform to the spatial dynamics required by the classical model. For instance, *TSP* affects tentacle patterning but not head or axis specification, while *HAS-7* appears to fine-tune local patterning rather than acting as a global inhibitory signal.

In light of these challenges, it seems increasingly unlikely that *Hydra* contains a molecular inhibitor that fully matches the assumptions of the classical activator-inhibitor model. Rather than seeking molecular players to fit an idealised framework, we propose reversing the approach: using a model-based analysis to explore whether the known interactions between *HyDkk1/2/4-A*, *HyDkk1/2/4-C*, and Wnt/*β*-Catenin signalling are sufficient to explain spatial patterning in *Hydra*.

The remainder of this paper is structured as follows. In the Results section, we introduce the new Mutual Inhibition (MI) Model that captures key molecular interactions between Wnt and Dkk proteins in *Hydra*. We present the model formulation, including biological justification and mathematical structure, followed by a comparison with experimental data and simulations of classical and novel perturbation scenarios. Next, we perform an in-depth mechanistic analysis for a reduced one-dimensional version of the model, using both numerical and analytical techniques to investigate conditions for pattern formation, including bistability and Turing instability. We explore the model’s robustness to parameter variations and qualitative modifications. Technical details, extended model variants, and supporting analyses are provided in the Supporting Information.

## Results

### Mutual Inhibition (MI) Model

To explore pattern formation ability of the Wnt-Dkk signalling system, we propose a mechanistic model describing interactions of *β*-Catenin/Wnt and the two Dkk-molecules HyDkk1/2/4-A and HyDkk1/2/4-C in *Hydra*. An overview of the system components and their interactions derived from various experiments is given in Fig. 1 (simplified) and Fig. 2 A (in detail). The model is denoted as the *Mutual Inhibition (MI) Model* after its core feedback loops that stand behind the pattern forming mechanism. The aim of our approach is to determine if a specific structure of the Wnt-Dkk signalling network is sufficient to explain the formation of the body axis in *Hydra*. The choice of the model components and their interactions is motivated by their experimentally documented function during *Hydra* axis formation and regeneration [26–30].

**Fig 1.**
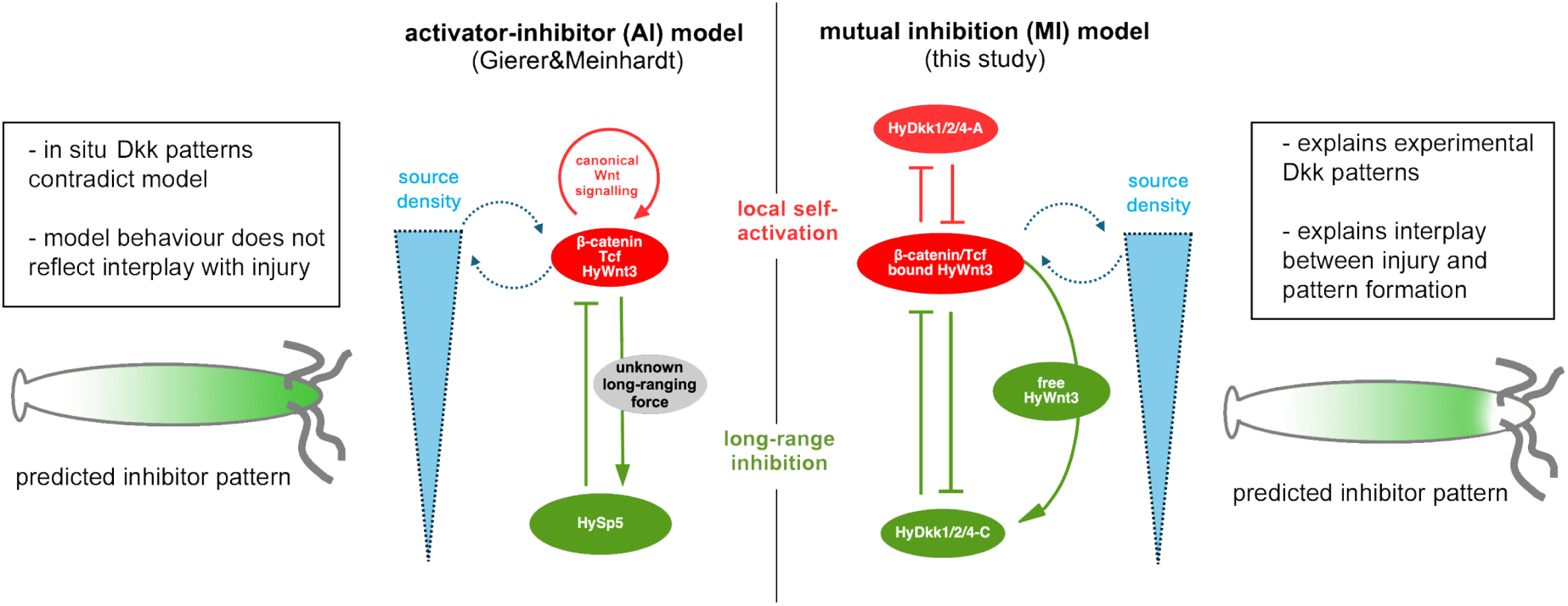
Schematic comparison of the activator–inhibitor (AI) model vs. the mutual inhibition (MI) model (this study). For clarity, coupling to the foot and tentacle patterning systems is not shown. In both models, a slow “source density” field (blue gradient) represents long-term positional information along the body axis, corresponding to the experimentally observed head-forming competence. Both models satisfy the general principle of local self-activation (red) and long-range inhibition (green). Grey ellipses indicate still hypothetical components or mechanisms. Model limitations and improvements are listed in the side boxes. Canonical HyWnt signalling is not explicitly resolved in the MI model for simplicity but may provide an additional or redundant mechanism for local self-activation. Predicted expression patterns of the inhibitors differ between the models (sketched polyps below). The mismatch between predicted and real HyDkk patterns led to neglect of Dkk molecules in the classical activator–inhibitor model.

**Fig 2.**
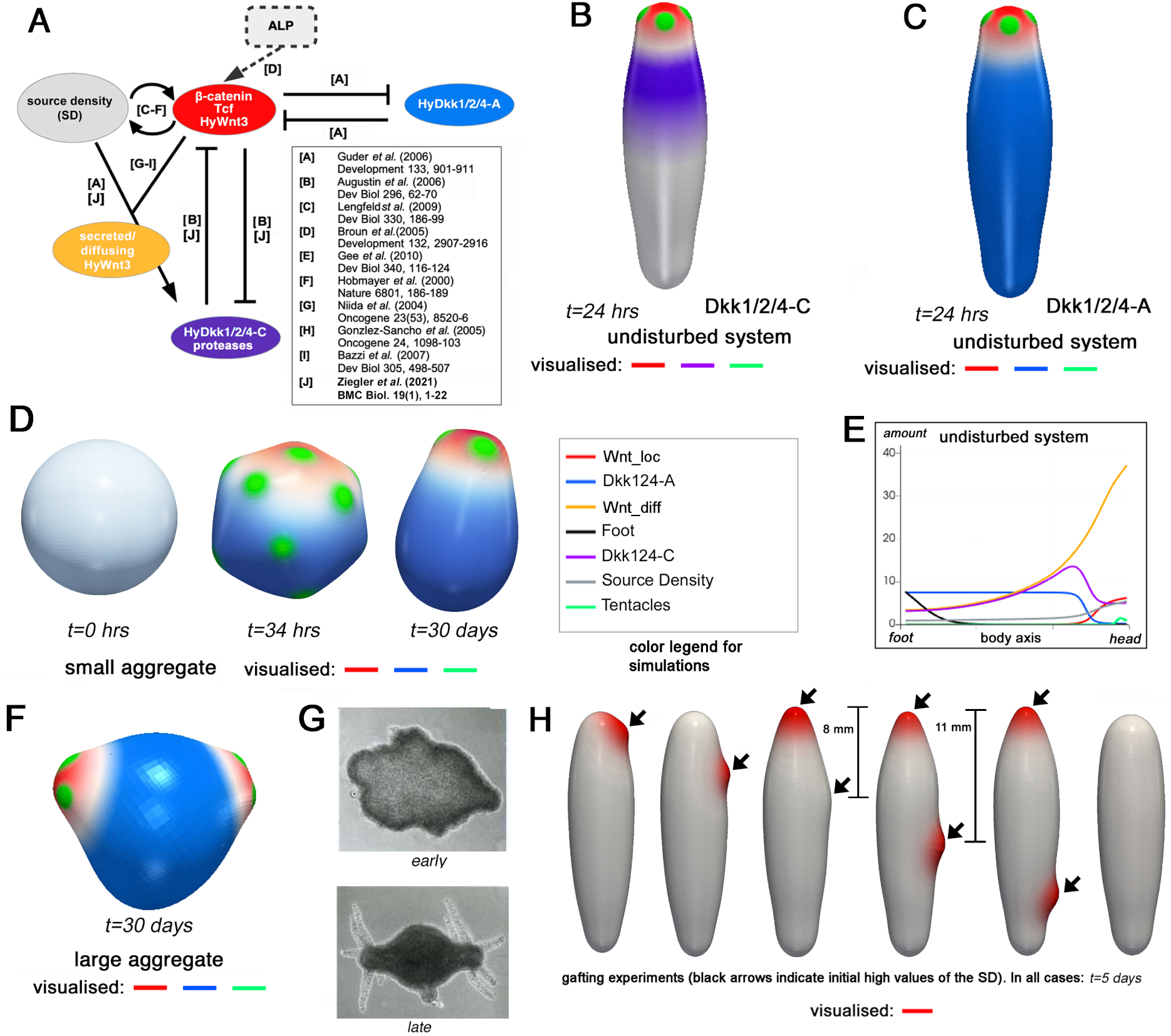
Overview, experiments, and simulations of HyWnt-Dkk-interactions and patterns. (A) Sketch of the *Hydra* body-axis formation network including main underlying references. For the sake of clarity, interactions with the tentacle and foot system are not shown. (B)-(C) Simulated activity fields corresponding to HyDkk1/2/4-C (B) and HyDkk1/2/4-A (C) in the undisturbed polyp. (D) Different stages of a simulated small aggregate; (E) concentration profiles of the different variables along the body axis in the undisturbed system; (F) late stage of a simulated large aggregate vs. early and late stages of experimental large aggregate (G). (H) simulated head formation resulting from virtual grafting experiments. Black arrows indicate initially given high values of the SD representing grafts. Colour-scaling is similar in all simulation snapshots in this figure (cf., color legend in the centre of the figure). Tentacle and foot structures are included for morphological realism only and do not affect the Wnt–Dkk-driven *de novo* patterning (see Fig. S 9).

### Model equations of the signalling system and their biological justification

The core of the model is given by reaction-diffusion-type equations describing dynamics of five components, see Eq.(1)-(5) and Table 1. The model variable [*W_l_*] represents the cell-local head-related molecules, e.g. reflected by the patterns of *HyWnt3, HyWnt9/10c, β-Catenin* and *Tcf* expression [26–28]. The model variable [*W_d_*] stands for the diffusing/free Wnt ligands (such as ligands of HyWnt3 or HyWnt9/10c) [36]. Variables [*A*] and [*C*] describe HyDkk1/2/4-A and HyDkk1/2/4-C, respectively [29, 30]. The model also incorporates the so-called source density (SD – variable [*S*]), a long-term store of information about the body axis gradient. Although its molecular identity remains unknown, its presence has been demonstrated by multiple experimental studies, see Refs. [4,9,15,37–40] and more details below. Our model representation is not meant to describe individual molecular species but to capture the effective regulatory activities of groups of molecules acting at similar spatial and functional levels. In this sense, [*W_l_*] and [*W_d_*] should be understood as coarse-grained quantities summarising the combined local and diffusible components of Wnt-related signalling, respectively. This abstraction reflects the high molecular redundancy and complexity of the *Hydra* pattern formation system and intends to capture system-level dynamics without explicitly resolving individual transcriptional or translational processes. The model equations read

**Table 1.** Model variables and their biological meanings.

| Variable name | Explanation |
| --- | --- |
| $[W_l]$ | cell-local head-related molecule complex comprising canonical Wnts, e.g. Hy-Wnt3 and HyWnt9/10c (e.g., Ref. [26–28]). |
| $[A]$ | Dickkopf1/2/4-A - a Wnt antagonist in *Hydra* [29, 30]. |
| $[W_d]$ | free (secreted/diffusing) canonical HyWnts (with patterns corresponding to nuclear β-Catenin activity) |
| $[C]$ | Dickkopf1/2/4-C - a Wnt antagonist in *Hydra* [30]. Optionally augmented by the astacin protease HAS-7 [35]. |
| $[S]$ | source density (SD) (also called e.g. *positional value* or *head forming competence*) - long-lasting information on the body-axis gradient; molecular/physical nature unknown. |

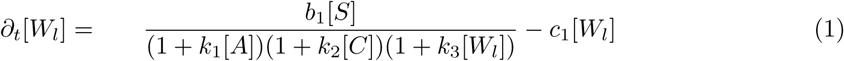

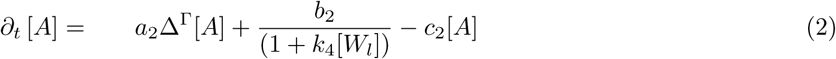

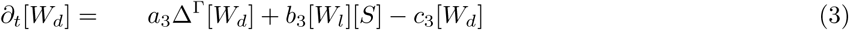

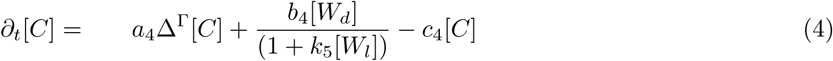

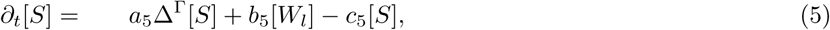

where Δ^Γ^[.] is the Laplace-Beltrami operator. The model is defined on a surface shell representing the Hydra tissue, and for the model analysis we additionally consider a reduced one-dimensional domain. The choice of the domain is discussed below. It should be noted that the model is not formulated as a mass-conserving two- or multi-compartment system. For example, the term *b*_3_[*W_l_*][*S*] in Eq. (3) represents the activation of diffusible Wnt-related activity by the local Wnt field and the source density, rather than a literal transfer of material. Accordingly, no corresponding influx term appears in Eq. (1), as [*W_l_*] acts as a regulatory source rather than a mass reservoir. Thus, formulations capture functional couplings between locally produced and diffusible components without implying mass conservation.

#### Cell-local Wnt dynamics

Eq. (1) describes the dynamics of the cell-local canonical Wnt activity, denoted as [*W_l_*], which summarises intracellular *β*-catenin/Tcf signalling and expression of canonical *HyWnt* genes. The first term describes production of [*W_l_*], which is promoted by the source density ([*S*]) (numerator). The first two inhibitory denominators represent the antagonistic action of Dkk1/2/4-A and Dkk1/2/4-C, consistent with experimental evidence for mutual repression between canonical Wnt and Dkk expression [29, 30]. The third denominator represents local self-limiting saturation, a natural assumption in the context of gene expression and protein synthesis. Finally, the last term in Eq. (1) accounts for degradation or turnover of the local Wnt activity. Together, these terms describe how positional competence (via [*S*]) and inhibitory feedback (via Dkks) jointly control the amplitude and spatial extent of local canonical Wnt signalling. Further supporting experimental evidence is given in [26–28, 36, 41, 42].

#### Dkk1/2/4-A dynamics

Eq. (2) describes the dynamics of HyDkk1/2/4-A ([*A*]), which acts as an inhibitor of canonical Wnt signalling. The first term represents (small) diffusion of [*A*] along the tissue surface, enabling spatial coupling between neighbouring cells. The second term describes production of [*A*], which occurs at a basal rate but is repressed by the local Wnt activity ([*W_l_*]), consistent with experimental observations showing mutual inhibition between HyDkk1/2/4-A and canonical Wnt signalling [29, 30]. The last term represents degradation or turnover of [*A*]. Together, these terms capture the role of HyDkk1/2/4-A as a short-range inhibitor that restricts and shapes the domain of local Wnt activation. Supporting experimental evidence is given in [29, 30, 35].

#### Diffusible Wnt dynamics

Eq. (3) describes the dynamics of diffusible Wnt activity, denoted as [*W_d_*], which represents the extracellular pool of secreted Wnt ligands acting over longer distances. The first term represents diffusion of [*W_d_*] along the tissue surface, modelling the lateral spread of secreted Wnt molecules. The second term describes the production of diffusible Wnt, which depends on both the source density ([*S*] – cf., below for more details) and the local Wnt activity ([*W_l_*]), expressed as the multiplicative term [*S*][*W_l_*]. This reflects that Wnt secretion requires two concurrent conditions: (i) sufficient tissue competence, encoded by [*S*], and (ii) active *β*-catenin/Tcf signalling, represented by [*W_l_*]. The choice of this coupling is biologically motivated by the observation that *β*-catenin/Tcf alone, while regulating many genes [43], may not be sufficient to trigger Wnt expression or secretion. Effective Wnt production likely requires additional, slower regulatory layers that determine whether a tissue region is permissive for head-related gene expression, such as stable epigenetic states, chromatin accessibility, or other long-term determinants of cellular identity. [*S*] thus represents a coarse-grained measure of these permissive conditions (cf., also below), while [*W_l_*] provides the transcriptional drive. Only where both factors are high can secreted Wnts be generated, ensuring spatial confinement of the diffusible Wnt source and robust axis polarity. The last term accounts for degradation or clearance of diffusible Wnt molecules. Together, these terms describe the production, spatial spreading, and turnover of secreted Wnt ligands that mediate longer-range feedback and contribute to axis stabilization. Supporting experimental evidence is given in [26, 28, 36, 41, 42]. Note that the actual diffusion coefficients of Wnt-related molecules in *Hydra* remain experimentally unconstrained; assuming moderate to strong diffusivity therefore represents a plausible but as yet unverified simplification. However, simulation shows that the patterns are robust to changes in the diffusion strength of the Wnt complex in our model (Supplementary Figure S 7 A-D).

#### Dkk1/2/4-C dynamics

Eq. (4) describes the dynamics of HyDkk1/2/4-C ([*C*]), a second Wnt antagonist with distinct regulatory control compared to [*A*]. The first term represents (small) diffusion of [*C*] along the tissue surface, allowing the molecule to act beyond the cells in which it is produced. The second term describes production of [*C*], which is positively regulated by diffusible Wnt activity ([*W_d_*]) and repressed by local Wnt activity ([*W_l_*]). This dual regulation reflects experimental findings showing that [*C*] expression increases in regions exposed to secreted Wnt ligands, but is downregulated in the head region where canonical Wnt/*β*-catenin activity is high [30]. The last term represents degradation or turnover of [*C*]. Together, these terms define [*C*] as an inhibitory component that is activated by long-range Wnt signalling but excluded from the head organizer itself, thereby reinforcing the spatial restriction of canonical Wnt signalling and contributing to the stability of the body-axis polarity. Supporting experimental evidence is given in [30, 35].

#### Source density dynamics

Eq. (5) describes the dynamics of the source density ([*S*]) encoding long-term positional information along the oral–aboral axis of *Hydra*. The first term represents diffusion of [*S*], modelling the slow redistribution of positional information through local interactions between neighbouring cells. The second term describes production of [*S*], which depends on local Wnt activity ([*W_l_*]) and captures the experimentally observed induction of long-lasting tissue competence by sustained *β*-catenin/Tcf signalling [19]. The third term accounts for decay, reflecting the gradual relaxation of positional information back to baseline levels over several days, consistent with regeneration and grafting experiments [27,37,44]. Together, these terms describe how transient Wnt/*β*-catenin signalling can establish as a long-term positional competence. Conceptually, [*S*] may represent an emergent property of the tissue, arising from a combination of molecular, cellular, or epigenetic processes, that defines whether a region is permissive for head-related gene expression and organiser formation. This abstraction allows [*S*] to capture the experimentally observed stability and spatial organisation of head-forming capacity without assuming a specific molecular identity. The source density evolves on a slower timescale than the Wnt–Dkk signalling system, reflecting its interpretation as a long-term positional memory field. This separation of timescales is implemented in the parameterisation of the model (cf., ‘Parameters and initial conditions’ section in the Supporting Information): the production- and decay rates of [*S*] are one to two orders of magnitude smaller than those of the Wnt and Dkk variables, and its diffusion coefficient is roughly three orders of magnitude smaller than that of [*W_d_*]. Supporting experimental evidence is given in [17, 27, 35, 37, 42, 44].

For more detailed experimental motivation and references supporting each term of the model equations, see Supporting Information, Section ‘Experimental and conceptual background of model components’.

### Model of the *Hydra* tissue

As the focus of this work is on the ability of the Wnt-Dkk signalling system to control the formation of a stable body axis, we investigate the proposed model both on a simplified one-dimensional domain representing the body axis (’1D model’) and in a more realistic geometry of the *Hydra* tissue (’pseudo-3D model’). For the latter, we adopt a mechano-chemical modelling framework coupling the signalling system with a model of an infinitely thin deforming tissue [45–47]. In the pseudo 3D model, the tissue surface is represented as an elastic shell governed by a Helfrich-type bending energy, with the local spontaneous curvature depending on the chemical fields (mainly [*W_l_*] and [*S*]). This coupling allows chemical patterning to influence the evolving tissue shape. Based on minimisation of the free energy describing elastic tissue deformations, the model results in a 4th order partial differential equation of the tissue evolution governed by the gene expression patterns resulting from the MI model. A detailed description of the physical assumptions, initial conditions, and chemo–mechanical coupling is provided in the Supporting Information (section “Mathematical framework for modelling pseudo 3D geometry”). In contrast to the fully coupled mechano-chemical models of [46–48], the current model does not account for any feedback from the mechanical properties of the tissue to the gene expression processes. Consequently, the pattern formation process is induced solely by the chemical signalling system. The purpose of including a realistic geometry in the evolving domain was to examine the potential impact of the underlying geometry on the sensitivity of pattern formation dynamics. To facilitate model analysis, we simplified the system to a one-dimensional domain with zero-flux boundary conditions, which serves as a simplified representation of the *Hydra* body column. The reduction in complexity enabled the efficient execution of numerical simulations and facilitated a more tractable analysis of the pattern formation mechanism.

### Model extensions to account for foot and tentacle dynamics

An additional version of the model includes two separate pattern-formation systems controlling foot and tentacle formation. These processes are not the focus of the present study, as they do not contribute to the pattern formation mechanism of the body axis. Foot and tentacle structures are included solely for visual and biological realism of the simulated *Hydra* morphology and to allow comparison with experimentally observed phenotypes, such as ectopic tentacles after ALP treatment. Both systems are represented by downstream activator–inhibitor modules that do not chemically feed back into the Wnt–Dkk mechanism responsible for axis pattern formation and might influence Wnt/Dkk only, if at all, indirectly via local surface deformations. To test this, we repeated the aggregate simulation shown in Fig. 2D with both the foot and tentacle modules removed (Fig. S 9). The resulting *de novo* symmetry breaking and final axis pattern remained unchanged, confirming that foot and tentacle formation have no influence on the Wnt–Dkk patterning mechanism. Further details of the foot and tentacle systems, including experimental justification of the model, are provided in the Supporting Information and in the references shown in Fig. 2.

#### Tentacle system

The tentacle subsystem is represented by an activator–inhibitor module that receives positive input from the source density ([*S*]) and negative input from local Wnt activity ([*W_l_*]). This captures experimental observations that tentacle primordia form preferentially in regions of intermediate head-forming competence, where [*S*] is high but *β*-catenin/Wnt locally suppresses this system [49]. The resulting field describes the periodic activation of tentacle-specific genes along the upper body column, consistent with observed tentacle spacing and regenerative behaviour.

#### Foot system

The foot subsystem is modelled analogously as an activator–inhibitor pair regulated by [*S*], but independent of the Wnt–Dkk head system. It represents the basal organiser region, stabilising the aboral pole. The foot activator is enhanced in areas of low [*S*], supporting the formation of a robust basal identity. Both subsystems are mathematically formulated in the Supporting Information.

### Model–Experiment Comparison and Validation

We evaluate the model’s ability to replicate key experiments by presenting the outcomes of various simulations and comparing them to experimental results. Most of the available molecular data on *Hydra* axis formation come from previously published *in situ* hybridisation experiments, which report spatial mRNA expression domains of individual genes. In contrast, the variables in our model represent coarse-grained activity fields of the underlying signalling pathways, combining contributions from several mRNA and protein species acting at similar spatial and functional levels. For example, [*W_l_*] reflects local canonical Wnt/*β*-catenin-related activity (as indicated by the expression patterns of *HyWnt3, HyWnt9/10c, β-catenin* and *Tcf*), whereas [*W_d_*] represents diffusible Wnt-related activity, for which no direct visualization is currently available in *Hydra*. In the following, the published *in situ* expression patterns are thus used as qualitative proxies for the spatial domains of the corresponding model activity fields. This correspondence is expected to be strongest for components with low effective diffusion, such as [*W_l_*] and the Dkk molecules, whereas it becomes less direct for diffusible components like [*W_d_*], for which no direct experimental visualisation is currently available.

### The MI model provides an explanation for the Dkk patterns that were not reproduced by previous models

We start with numerical analysis of the wild-type pattern that resembles experimental observations (Fig. 2 B-C and Ref. [29, 30]). Such pattern can be established *de novo* if the initial conditions provide a sufficiently strong canonical Wnt signal localised at the head end, what corresponds to the head cutting experiments with a localised signal stemming from the injury. With respect to the resulting stable patterns, the simulations predict the lack of both HyDkk1/2/4-related molecules in the hypostome, showing a sharp expression border beneath the tentacles, *HyDkk1/2/4-C* expression fading out in the aboral direction (Fig. 2 B and E), and *HyDkk1/2/4-A* strongly expressed in the entire body column (Fig. 2 C,E). The simulated [*W_l_*] activity field is restricted to the head region, fading within and below the tentacle zone (Fig. 2 B,C,E). Also, the temporal evolution of canonical Wnt activity matches qualitatively and quantitatively well between experimental observations and our simulations (Supplementary Figure S 8 F vs. Ref. [26]).

The difference in the two HyDkk patterns results from the qualitative differences in the corresponding production terms in the model equations. While HyDkk1/2/4-A is assumed to be constantly produced in the absence of repressing signals, HyDkk1/2/4-C production is modelled downstream of the HyWnt signalling and thus fades out in aboral direction. To explore the role of the constant production term, we additionally simulated the system with the HyDkk1/2/4-A production depending on SD instead of considering a constant expression. It led to a graded *HyDkk1/2/4-A* expression along the body axis fading out in ab-oral direction (Supplementary Figure S 7 I–J). In this modified system, the expression of [*W_l_*] appeared to be distinctly increased in the budding zone (compared to the original model), indicating that body-wide expression of *HyDkk1/2/4-A* might be involved in controlling/suppressing bud formation.

It is important to notice that although both Dkk molecules are produced in overlapping domains and are based on mutual inhibition loops with canonical Wnt signalling, HyDkk1/2/4-A can be seen as a part of the local activation, whereas HyDkk1/2/4-C is rather involved in long-range inhibition (c.f., Fig. 1, Fig. 2, and Supplementary Fig. S 7 E–H). This apparent contradiction between antagonistic activity and similar expression patterns arises from the fact that the local activation is realised through an inhibition of the inhibitor HyDkk1/2/4-A (c.f., Fig. 1 and Fig. 2). Thus, the inverse expression pattern of HyDkk1/2/4-A is comparable to the activator pattern in the activator-inhibitor model and indeed, the inhibitory effect of HyDkk1/2/4-A on canonical Wnt signalling is required for pattern formation (Supplementary Figure S 8 E).

### The MI model can describe self-organisation

To explore whether the observed ability of self-organised axis formation critically depends on the initially provided geometry or chemical (SD) gradient, we simulated self-organisation in *Hydra* aggregates. The experiment was modelled assuming a symmetric tissue sphere and a random distribution of all considered biochemical components, including the SD. The simulations showed *de novo* pattern formation resembling the experimentally observed patterns in adult polyps (Fig. 2 D,F,G). Notably, small initial spheres evolved into a single polyp (Fig. 2D), while larger aggregates gave rise to multiple heads (Fig. 2F), in good agreement with the experimental observations (Fig. 2G). In conclusion, the MI model reproduces correctly the scaling of *de novo* pattern-formation system.

### The MI model requires a strong localised signal for head regeneration in cutting experiments

One of the key challenges to *Hydra* models of head regeneration is the recently demonstrated role of strong localised signalling in cutting experiments. These experiments show that removing the head without creating a wound does not lead to regeneration; only the activation of strong signalling and a localised increase in *Wnt3* expression as part of the wound response enables regeneration [19]. This finding is particularly significant as it directly challenges the classic *de novo* pattern formation mechanism described by activator-inhibitor models. In former theoretical work, we postulated the necessity of bi-stability in a model to account for such phenomena [50]. The MI model meets these conditions, describing the absence of pattern formation under certain initial conditions, specifically, when an area with high *Wnt3* expression and source density levels is removed, Fig. 2 H – right hand side. Conversely, the introduction of a strong signal, such as that triggered by the wound response, facilitates pattern formation. A systematic analysis of this mechanism is provided in the next section.

### The MI model reproduces the transplantation experiments

Further, we simulate transplantation experiments (Fig. 2 H) motivated by the seminal experiments presented in Ref. [37, 51, 52]. These experiments showed very early on that, under certain circumstances, tissue fragments transplanted from one polyp to the other can lead to the growth of a secondary body axis. Our simulations reproduce key properties. Head self-inhibition results in suppression of the head regeneration if two initiating signals are too close (3th simulation snapshot in Fig. 2 H). Moreover, without the head at the oral pole, grafts can induce a secondary axis at the arbitrary locations (1th and 2th simulation snapshot in Fig. 2 H). Simulations showed emergence of the secondary head in virtual graftings only when the distance is above approx. 50 % of the body length, which agrees with quantitative experimental observations [42] (4th and 5th simulation snapshot in Fig. 2 H). All these features of the *Hydra* patterning system stay in agreement with experimental observations [37, 42, 52, 53].

### Further *in silico* experiments

In the next step, we verify the role of different model components by simulating their manipulated levels and comparing the resulting patterns to experimental data. First, we examine how system performance depends on the presence of the two HyDkk1/2/4-related molecules. Specifically, we simulate the undisturbed polyp system until the head pattern is established, as shown in Fig. 2 B,C and 3 A (left-hand side). We then perturb the system by virtually removing the expression of one of the two *HyDkks*. In line with experimental evidence, we observe a distinct expansion of [*W_l_*] in the head region following the removal of HyDkk1/2/4-A (Fig. 3 A (middle) and Ref. [29]) and body-wide ectopic activation of the [*W_l_*] field after removing *HyDkk1/2/4-C* (Fig. 3 A, right-hand side as well as Ref. [30]). The only difference between the simulations and experiments is that [*W_l_*] production appears relatively diffuse and homogeneous after reducing the number of *HyDkk1/2/4-C* -expressing cells (Fig. 3 A, third snapshot), whereas experiments report a more spotty pattern of *HyWnt3* expression [30]. This discrepancy is likely due to the fact that, as mentioned earlier, the model does not distinguish between the cell-local HyWnt3 and *β*-catenin or Tcf, whereas in reality, these factors may belong to two distinct pattern-formation systems operating at different spatial scales [54]. However, for the sake of model simplicity, further distinctions between *β*-catenin-driven axis formation and HyWnt3-driven head/organisation formation (cf. [54]) are not considered in this study, as the primary focus is on the general principles of HyWnt-Dkk-driven axis formation.

**Fig 3.**
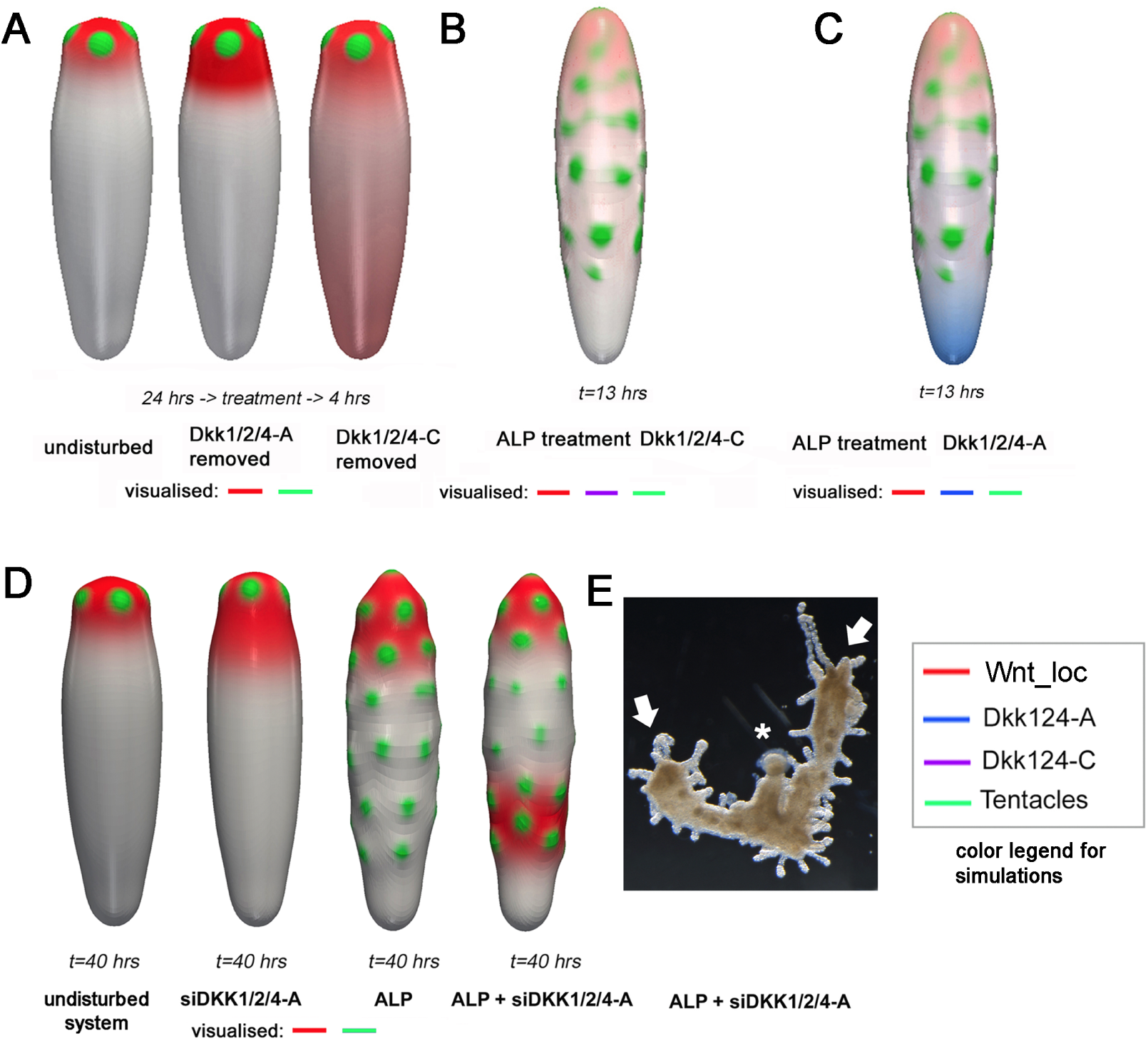
Simulations and experiments of manipulated HyWnt and HyDkk-levels. (A) simulation snapshots of the undisturbed polyp vs. virtual removal of *HyDkk1/2/4-A* vs. *HyDkk1/2/4-C* expression showing the [*W_l_*] activity field (corresponding to *β*-Catenin/Tcf/bound HyWnt3 domains) and tentacles. (B) Simulations of HyDkk1/2/4-A vs. *HyDkk1/2/4-C* expression after ALP-treatment. (D) Simulation snapshots of [*W_l_*] and tentacle patterns of the undisturbed system vs. different combinations of ALP-treatment and *HyDkk1/2/4-A* knockdown. (E) experimental picture of double axis formation after ALP + siDkk1/2/4-A treatment, arrows indicate axes, asterisk represents foot. Colour-scaling is similar in all simulation snapshots in this figure (c.f., colour legend on the bottom right-hand side).

In addition, we simulate the activation of canonical HyWnt signalling via ALP treatment (Fig. 3 B,C, as well as Supplementary Figure S 8 A–D). Experimentally, both HyDkks are suppressed by this treatment, along with the development of ectopic tentacles along the body column [29,30]. However, *HyDkk1/2/4-C* expression appears to be more sensitive to *β*-catenin/HyWnt3 levels, as the reduction in *HyDkk1/2/4-A* expression occurs later than the reduction in *HyDkk1/2/4-C* expression [29, 30]. These observations align with our simulation results during ectopic tentacle development (Fig. 3 B,C). However, we note that in later stages of the simulations, Dickkopf patterns re-establish (1D concentration profiles in Supplementary Figure S 8 A–D), suggesting that the model can reproduce the data only as transient patterns. This may suggest that ALP-driven effects are initially present but reversible. Finally, we simulate recent *HyDkk1/2/4-A* knockdown experiments [35] with and without ALP treatment (Fig. 3 D). Similar to the experimental results (Fig. 3 E), the combination of ALP treatment and *HyDkk1/2/4-A* knockdown leads to the development of a secondary ectopic axis, marked by an additional region with high activity of the canonical HyWnt signalling field in our simulation results, but again, this is observed as a transient pattern.

### Analysis of the Mechanism Underlying Pattern Formation

To determine under which conditions the MI system can exhibit symmetry breaking and pattern formation, we analyse the structure and stability of spatially homogeneous steady states. Stability of these states indicates that no patterns form in their vicinity, whereas a loss of stability, triggered by sufficiently contrasting diffusion rates, can lead to the emergence of well-known Turing patterns. The underlying mechanism, called the diffusion-driven instability (Turing instability) is a local bifurcation from a spatially homogeneous steady state due to a sufficient space scale separation in diffusion rates. Systems coupling diffusive and non-diffusive components can, additionally, exhibit *far-from-equilibrium* patterns characterised by jump-discontinuity [55–57]. Such solutions emerge due to the system’s bistability. While a Turing instability may act as a trigger, the emergence of *far-from-equilibrium* patterns can occur independently of the Turing mechanism. Models exhibiting both bistability and DDI have been studied both, in full reaction–diffusion systems [58] and in settings with non-diffusing model components [56]. For linearised and nonlinear stability analysis theory for reaction-diffusion-ODE models we refer to [55, 59, 60].

To better understand the nature of the patterns observed in our simulations, we analyse the stability of branching stationary solutions near the DDI bifurcation point. Since a rigorous analysis of complex models is often infeasible, we complement it with numerical analysis. Guided by analytical insights, we analyse the model for various fixed parameter values. This is further supported by a sensitivity analysis, which assesses the model’s robustness to parameter and nonlinearity variations and confirms the validity of the results within specific parameter regimes.

### One-dimensional model reduction

In this section, we focus on a one-dimensional version of the model given by equation (1)-(5) to systematically analyse its dynamics as a function of the parameters and initial conditions. Our goal is to provide a deeper understanding of the simulation results discussed earlier. We begin by examining the model’s ability to generate patterns. This involves analysing the mathematical structure of the model, the ability of symmetry breaking and the stability of the emerging patterns.

The reduced model is obtained by approximating the complex domain of *Hydra* by an interval [0, 1] and rescaling time and model variables in equations (1)-(5). The technical details of the model reduction, analytical approach, and numerical implementation are provided in the Supporting Information. To compare the reduced system with the original full model, we apply a computational fitting procedure using data derived from the 3D simulation (Fig. 2 E). These data are obtained by averaging along the axis perpendicular to the body axis, followed by min–max normalisation. Parameter estimates are identified by minimising the least-squares residual between the data profiles and the output of the reduced model. This procedure demonstrates that the reduced model reproduces the same dynamics as the original, with appropriate parameter adjustments. Further numerical details are included in the Supporting Information Table S 1.

### The MI model reveals bistable behaviour in uniform steady-state structures

Analysis of the structure of the spatially uniform steady states demonstrates a bifurcation from a semi-trivial state ([*W_l_*] = 0, [*A*] = *β*_1_, [*W_d_*] = 0, [*C*] = 0, [*S*] = 0), where *β*_1_ = *b*_2_*/c*_2_, resulting in the existence of additional spatially homogeneous solutions and a change of stability, see Figure 4. We identify two complementary bifurcation parameters, *β*_1_: = *b*_2_*/c*_2_ and *β*_6_: = *b*_1_*/*(*c*_1_*c*_5_), which describe the effective production rates of [*A*] and [*W_l_*], respectively. More precisely, linear stability analysis of the semi-trivial steady state yields an instability condition, *β*_6_ *>* 1 + *β*_1_, which marks the bifurcation point and highlights the complementary relationship between the two bifurcation parameters. The corresponding bifurcation diagrams are shown in Fig. 4 A and Fig. 4 B. The remaining parameter values are fixed according to the parameter fit to the three-dimensional model discussed above (see Supplementary Table S 1).

**Fig 4.**
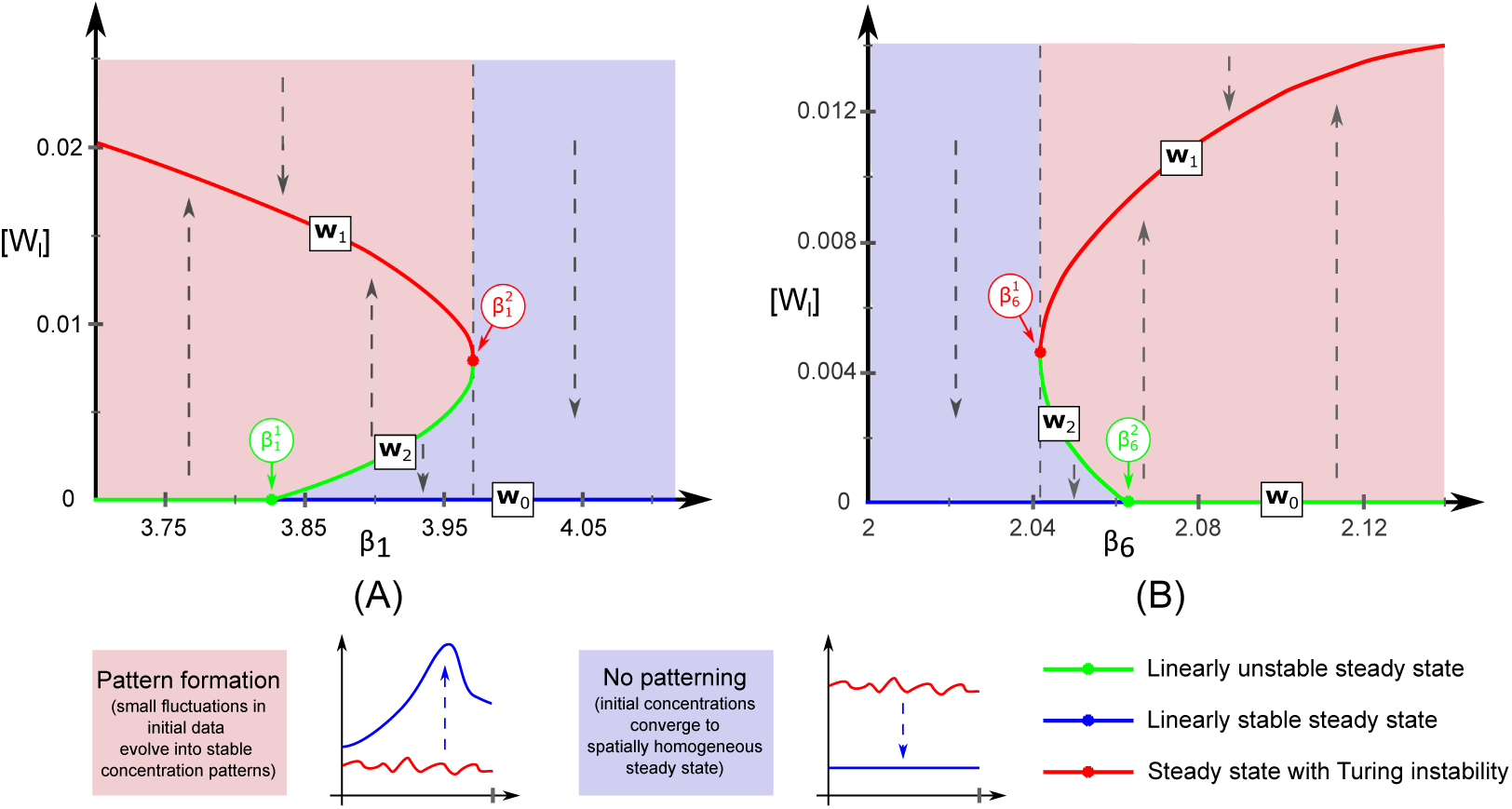
Bifurcation diagram illustrating the qualitative changes in pattern formation in the one-dimensional version of model (1)–(5), induced by variations in the rescaled reaction parameters *β*_1_: = *b*_2_*/c*_2_ (effective production rate of [*A*]) and *β*_6_: = *b*_1_*/*(*c*_1_*c*_5_) (effective production rate of [*W_l_*]). The vertical axis represents the first component of the homogeneous steady states. Arrows indicate whether initial conditions converge to the pattern formation regime or decay towards the trivial steady state. Coloured bullets mark the boundaries of the bistable region: 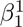 and 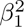 in panel (A); 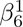 and 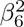 in panel (B). All other model parameters are set to the values listed in Supplementary Tab. S 1.

The structure of the bifurcation diagram and the stability of different branches of spatially uniform solutions are determined based on the model analysis presented in the Supplementary Information (Lemma S1-S4 and Corollary S1) and numerical computations, employed whenever a complete analytical understanding was not feasible. To this end, we investigate the model for varying either *β*_1_ or *β*_6_, while fixing all other parameters. As theoretically predicted, for sufficiently small values of *β*_1_, we observe instability of the semi-trivial stationary solution, denoted by ***w***_0_, and existence of exactly one non-trivial spatially uniform steady state ***w***_1_ which is stable for the system without diffusion. Numerical computation of the system linearised at this positive steady state indicates Turing instability. This stays in agreement with the model simulations showing globally stable pattern formation. Above the critical value (transcritical bifurcation point), the semi-trivial state becomes stable and gives branching to an unstable homogeneous steady state ***w***_2_ to eventually collide with ***w***_1_ for the parameter value corresponding to the saddle-node bifurcation point. This results in a bistable regime for the parameter 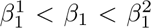. This means that for initial concentration values in the basin of attraction of the trivial steady state ***w***_0_, there is no pattern formation, and initial data decay to the trivial steady state ***w***_0_. On the other hand, if initial data lie in the basin of attraction of the steady state ***w***_1_, then we observe the formation of patterns. Here, unstable steady state ***w***_2_ acts like a separation hyperplane between two regimes, see Figure 5. For 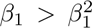 no pattern formation occurs and the only existing spatially uniform steady state ***w***_0_ becomes a stable attractor. Analogous results are obtained with respect to the bifurcation parameter *β*_6_, as shown in the bifurcation diagram Fig. 4 B.

**Fig 5.**
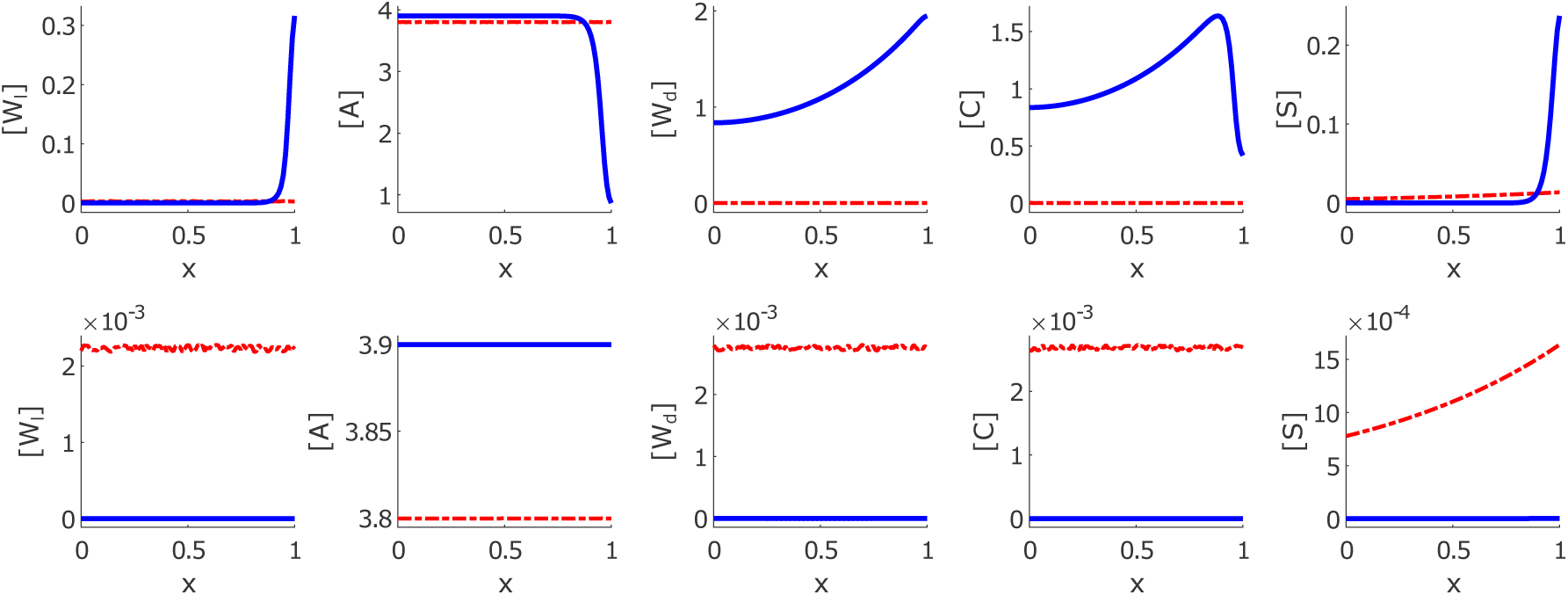
Example of bistable behaviour in the one-dimensional reduced model. Initial data are taken as small perturbations of the homogeneous steady state ***w***_1_. When small perturbations lie in the basin of attraction of ***w***_1_, pattern formation occurs and initial data evolves to the steady-state patterns (first row). Otherwise, when the steady state is perturbed in the *direction* of the trivial steady state ***w***_0_, no pattern formation occurs and initial concentrations decay to the trivial steady state (second row). Parameter values used in the simulations are given in Supplementary Table S 1. Blue colour shows the final concentration profiles, while red colour denotes the initial concentration values.

Translated into the language of experimental research, this means that pattern formation or regeneration does not always occur as predicted by the classical activator-inhibitor model, but can critically depend on the initial conditions. This, in turn, explains why the MI model aligns with the experimental observation that regeneration only happens when the initial conditions provide a sufficiently strong stimulus through wound signalling.

### MI model exhibits Turing pattern formation

The Turing instability leads to spatially heterogeneous solutions that may converge to stationary patterns. To gain a deeper understanding of the emerging patterns, particularly in the bistable region, we conduct a bifurcation analysis of branching solutions near the onset of Turing instability, see Supporting Information for details. Since not all required expressions can be derived explicitly, we compute certain terms numerically. We examine two sets of model parameters to encompass both regimes—one with hysteresis and one without—and use the largest diffusion coefficient as the bifurcation parameter. We then numerically compute the critical value and apply classical bifurcation theory to construct asymptotic expressions for branching stationary solutions and assess their linear stability. In both cases, our analysis indicates that the branching solutions remain stable, consistent with numerical simulations for the corresponding parameter values. Although our approach does not constitute a rigorous mathematical proof, it provides strong evidence supporting the presence of Turing patterns.

### MI model is robust with respect to parameters’ perturbations

Next, we numerically investigate the robustness of pattern formation with respect to variations in model parameters. To demonstrate this robustness, we use the parameter values listed in Supplementary Table S 1. We then explore the hypercube in parameter space for rescaled reaction rates *β_i_* within a large range [*β_i_/*100, 100 *β_i_*], parameters are therefore varied by a factor of 10,000. To estimate the regions within this hypercube where pattern formation occurs, we employ Markov Chain Monte Carlo (MCMC) methods. The criterion for admissibility is the existence of at least one homogeneous steady state exhibiting Turing instability. For the chosen parameter values, we did not identify any cases where this criterion was not met, indicating the robustness of the observed pattern formation. Of course, the range of parameters studied here did not exceed the previously discussed points of transcritical bifurcation, beyond which the only stable equilibrium state of the system would be the semi-trivial state. Another critical parameter is, as discussed before, the largest diffusion in the system, which has already been discussed earlier and must be sufficiently large in order to achieve Turing instability.

### MI model is robust with respect to a range of qualitative changes

Finally, we assess the robustness of the reduced model (for equations, cf. Supporting Information) with respect to modifications in the reaction terms. We consider two variations of the model. In the first, the reaction terms are multiplied by their respective denominator expressions. This modification can be interpreted as describing inhibition at the level of degradation or receptor binding, rather than at the level of production. Another system’s perturbation concerns using higher-order terms in Hill functions, i.e. modulating the inhibition process. For the parameter values examined, both variations exhibit the same qualitative pattern formation behaviour as the original one-dimensional model, suggesting its robustness. For further technical details, see the Supporting Information.

## Discussion

Symmetry breaking and pattern formation in *Hydra* have been extensively studied through the activator-inhibitor model, which has provided significant insights into the underlying processes [17, 61], and identified the pattern formation principle of local activation and long-range inhibition (LALI) [13, 32]. However, discrepancies have emerged between predicted and observed patterns, highlighting the limitations of the classical activator-inhibitor framework. Main challenges are: (1) the elusive identification of inhibitors consistent with the theory [48]; (2) observed patterns of key Wnt inhibitors, such as Dkk, contradicting the activator-inhibitor model [29, 30]; (3) the failure of the model to predict the necessity of wound healing signals for regeneration [19]; and (4) the requirement for specific nonlinear kinetics, which are difficult to justify from molecular or experimental perspectives [13, 62, 63].

To address these challenges, we developed the Mutual Inhibition (MI) Model, which: (1) is based on known Dickkopf inhibitors and their experimentally determined interactions; (2) generates patterns that align with experimental data for all required molecules, including both Dickkopfs; (3) produces spatial patterns only when initial conditions are sufficiently far from equilibrium, consistent with wound signalling; and (4) remains robust to parameter disturbances and qualitative changes. The MI model was validated through its alignment with a broad range of experimental data and manipulations, including genetic and pharmacological treatments and classical grafting experiments. Additional control simulations demonstrated that the body-axis patterning in the model originates from the core Wnt–Dkk interactions and does not depend on the presence of the auxiliary foot and tentacle systems (Fig. S 9).

The Mutual Inhibition (MI) model presented here is best understood as a biologically grounded realisation of the general local–activation/long–range–inhibition (LALI) principle of *de novo* pattern formation, rather than as an alternative to it. In their original work, Gierer and Meinhardt already demonstrated that LALI can be implemented by different reaction–diffusion schemes, including both the classical activator–inhibitor system and the activator–depleted–substrate mechanism [8, 17]. Later, Meinhardt further generalised this concept to multi-component systems, emphasising that “activation” and “inhibition” can be properties of subsystems (for example realised by mutual inhibition or inhibition of an inhibitor) rather than of single molecular species [16, 17]). In this spirit, the MI model implements the LALI principle through the experimentally established antagonistic interactions between Wnt and Dkk molecules in *Hydra*. Short-range “activation” arises from the mutually inhibitory Wnt–Dkk subsystem, which stabilises local high Wnt activity, while long-range “inhibition” is mediated by diffusible components and their spatial spread (in particular [*W_d_*] together with the Dkk gradients) also comprising mutually inhibitory motifs. Thus, the MI model links the abstract LALI framework to specific molecular players and interaction motifs that account for the observed Wnt/Dkk patterns in *Hydra*. It is important to note that the mutual inhibition between [*W_l_*] and [*C*] is not required for the emergence of a stable body-axis pattern, as demonstrated in control simulations (Fig. S 7 E–H). The core mechanism of pattern formation arises from the overall Wnt–Dkk network architecture that realises the LALI principle.

The new model provides insights into how Dkk molecules and canonical Wnt signalling may lead to self-organised pattern formation in *Hydra*, specifically through repeated motifs of mutual inhibition. Although the Dkk–Wnt interplay is essential for various developmental processes [21, 64], the underlying patterning mechanisms have remained unclear. The MI mechanism presented here may represent a general and fundamental patterning motif, extending beyond the phylum Cnidaria. However, a sufficiently strong diffusion for (some) Wnt molecules is still a critical assumption in the presented MI model. While studies in other organisms indeed suggest such diffusion [65–67], direct evidence in *Hydra* is still lacking. The assumed diffusion may also be interpreted as an effective measure for various propagation mechanisms, including the synergistic spread of multiple Wnts [36], biomechanical cues [68], bio-electrical signalling [69], or active transport along cell protrusions [70].

Beyond the biological insights, the study proposes a paradigm shift in pattern formation research. Traditionally, the top-down approach dominated, where theoretical models guided the search for matching molecular components. However, as recent advancements show a plethora of two- and three-component networks capable of self-organisation into Turing-like patterns [63, 71], a top-down approach may be inefficient. The present study adopts a bottom-up approach, building a model based on experimentally identified system components and interactions. By iteratively combining this approach with theoretical studies, we can develop effective models and generate specific questions for hypothesis-driven experimental research.

Additionally, the present study does not distinguish between body axis and head formation in *Hydra*, although recent findings suggest that these processes should be analysed separately [54]. The MI mechanism is more closely associated with the formation of the body axis, as the observed patterns establish on a large (body axis) scale rather than on the smaller scale of the head organizer. This suggests that other canonical Wnts, such as *HyWnt9/10c*, may play a central role in the activation process, as indicated by its early expression during regeneration [36]. Indeed, a *HyWnt9/10c* knockdown results in complete regeneration failure, while *HyWnt3* knockdown still allows tentacle formation, indicating that the body axis remains intact [19].

In the present study, various components such as *β*-Catenin, HyTcf, *HyWnt9/10c*, and *HyWnt3* were grouped into one molecular class, further distinguishing between cell-local and long-range components. This categorisation highlights several gaps, such as the broader spatial domain of *β*-catenin’s nuclear activity compared to *Wnt3* expression. This suggests that canonical Wnt signalling may involve an additional mechanism to restrict *Wnt3* expression patterns [54]. A more detailed view of these components in interaction with Dkk molecules in *Hydra* is needed to better distinguish between head and body axis formation. Further research could investigate the effects of *Dkk* overexpression or knockdown on the distribution and levels of these Wnt molecules.

In recent years, several complementary frameworks have been proposed to explain pattern formation in *Hydra* beyond purely chemical interactions. Braun and Agam highlighted the role of global mechanical feedbacks and stochastic morphogenetic fluctuations, describing regeneration as a canalised but noisy process governed by interactions between tissue tension, geometry, and biochemical activity [72–74]. In parallel, groups of Keren, Roux, Tomancak and co-workers revealed that the supracellular actin fiber network forms an active nematic field whose topological defects coincide with organiser sites, providing a mechanical mechanism for head and foot initiation [75–78]. These studies demonstrated that anisotropic stretch and curvature can bias defect orientation and thus organiser positioning, suggesting that cytoskeletal alignment and tissue mechanics act as self-organising pre-patterns for morphogenesis. Finally, mechanochemical theoretical and experimental approaches developed recently [47, 48] have explicitly linked Wnt-related morphogen regulation with tissue deformation, showing that chemical and mechanical feedbacks together can stabilise or reshape axis formation and induce *de novo* pattern formation.

Our current MI model focuses on biochemical patterning at the body-axis scale, describing how mutual inhibition between Wnt and Dkk molecules can self-organise stable polarity and long-term positional information. This distinguishes the MI mechanism from recently proposed mechanical or mechanochemical frameworks, which primarily address the emergence of local head organizers.

Hence, axis formation and organizer formation may represent distinct, yet coupled processes that operate on different spatial and temporal scales [54]. Alternatively, the chemical and mechanical mechanisms may act in parallel or redundantly, contributing to the robustness of pattern formation in regenerating tissue. Finally, all three layers —biochemical feedback, cytoskeletal alignment, and tissue mechanics— could be parts of an integrated system in which mechanical and topological cues modulate Wnt–Dkk signalling and *vice versa*. Clarifying how these processes interact to coordinate axis and organiser formation represents an important direction for future research.

In summary, over the last decade, our understanding of *Hydra* pattern formation has advanced considerably. Early models provided a basic framework but did not capture the full complexity of pattern formation. The identification of distinct patterning loops, along with recognition of non-chemical processes such as cytoskeletal organisation and bioelectrical signalling, has added additional layers of complexity [48, 69, 79–83]. Moreover, the role of Wnt antagonists in regulating pattern formation has revealed a far more intricate picture than initially understood [29, 30, 33, 35]. The MI model presented here has allowed for a better exploration of different aspects of Wnt signalling that indicated gaps in the current systematic understanding of pattern formation in *Hydra*. The next challenge will be to integrate these findings into new experimental designs to achieve a coherent understanding of developmental patterning.

The MI model is also of significant theoretical interest, as it exemplifies a system that couples diffusing and cell-local components. Such reaction-diffusion-ODE models have been the focus of analytical research in recent years, as they can exhibit dynamics that extend beyond classical Turing systems [55, 84–88]. Notably, in models where cell-local components interact with a single diffusing morphogen, all Turing patterns may become unstable [59, 89], giving rise to *far-from-equilibrium* patterns characterised by jump discontinuities [56, 57, 90]. However, models that couple space-dependent ODEs with multiple diffusive components, such as the MI model, remain largely unexplored. Our findings demonstrate that in such models, stable Turing patterns can still emerge, even in conjunction with bistability, highlighting the need for further theoretical investigation.

## Acknowledgments

This work was supported by the Deutsche Forschungsgemeinschaft (DFG) through Germany’s Excellence Strategy EXC-2181/1 – 390900948 (Heidelberg STRUCTURES Excellence Cluster) and through SFB1324 (projects B05 to AM-C and MM, B07 to SÖ, and A05 to TWH and AT). The funders had no role in study design, data collection and analysis, decision to publish, or preparation of the manuscript. AM-C, MM, SÖ, TWH and AT received salary support from the above DFG funding programs. The authors would like to thank Szymon Cygan and Finn Münnich for valuable discussions on model analysis.

## Notes

### Competing Interest Statement

The authors have declared no competing interest.

### Summary of Updates

In this revised version of the manuscript, we have substantially improved clarity, structure, and interpretability in response to peer-review feedback. We clarified the functional roles of all model components, explicitly distinguished between cell-local and diffusible Wnt/Dkk activities, and strengthened the connection between model variables and experimental in situ data. The manuscript now provides a clearer explanation of how the Mutual Inhibition model relates to and extends classical LALI frameworks. We added new simulations demonstrating that tentacle and foot modules do not influence axis formation, expanded the analysis of bistability and diffusion-driven instability, and improved the mathematical presentation in both the main text and Supplementary Information. All figures, variable notations, and gene/protein nomenclature were updated for consistency with biological conventions. Overall, the revised manuscript provides a clearer, more rigorous, and more biologically grounded presentation of the Wnt-Dkk-based pattern formation mechanism in Hydra.

