## Supporting Information File for "Mutual Inhibition Model of pattern formation: The role of Wnt-Dickkopf interactions in driving *Hydra* body axis formation"

Moritz Mercker[1,*], Alexey Kazarnikov[1], Anja Tursch[2], Thomas Richter[3], Suat Özbek[2], Thomas Holstein[2], Anna Marciniak-Czochra[1]
**1 Institute for Mathematics and Interdisciplinary Center of Scientific Computing (IWR), Heidelberg University, Heidelberg, Germany**
**2 Centre for Organismal Studies (COS), Heidelberg University, Heidelberg, Germany**
**3 Institute of Analysis and Numerics, University Magdeburg, Magdeburg, Germany**

*****

#### One-dimensional model reduction

Before presenting the mathematical formulation, it is important to note that the model is defined on a functional rather than molecular level. Each variable represents the aggregate activity of several signaling components acting at similar spatial and functional scales, rather than a single gene or protein species. For instance, $[W_l]$ and $[W_d]$ denote local and diffusible Wnt-related activities, respectively, combining the contributions of multiple Wnt mRNAs and proteins as well as beta-catenin and Tcf. The Dickkopf components correspond to the activities of HyDkk1/2/4-A and HyDkk1/2/4-C. Consequently, the model equations describe effective regulatory interactions and are not intended to represent mass-conserving fluxes or stoichiometric relations between individual molecular species.

We consider one-dimensional spatial domain $\Omega = [0, 1]$. The system (1)-(5) (main manuscript) reads

$$\begin{aligned}
\partial_t[W_l] &= \frac{b_1[S]}{(1+k_1[A])(1+k_2[C])(1+k_3[W_l])} - c_1[W_l];\\
\partial_t\,[A] &= a_2\frac{\partial}{\partial x^2}[A] + \frac{b_2}{(1+k_4[W_l])} - c_2[A];\\
\partial_t[W_d] &= a_3\frac{\partial}{\partial x^2}[W_d] + b_3[W_l][S] - c_3[W_d];\\
\partial_t[C] &= a_4\frac{\partial}{\partial x^2}[C] + \frac{b_4[W_d]}{(1+k_5[W_l])} - c_4[C];\\
\partial_t[S] &= a_5\frac{\partial}{\partial x^2}[S] + b_5[W_l] - c_5[S],
\end{aligned} \tag{S1}$$

supplemented with initial conditions and homogeneous Neumann (zero-flux) boundary conditions for all diffusing components.

To streamline model analysis, we reduce the number of parameters by deriving the system's non-dimensionalised form. To this end, we rescale the variables

$$\alpha_5[W_l] = [\tilde{W_l}],\ k_1[A] = [\tilde{A}],\ \alpha_4 k_2[W_d] = [\tilde{W_d}],\ k_2[C] = [\tilde{C}],$$

where $\alpha_i = \dfrac{b_i}{c_i}$, $i = 1, \ldots, 5$, rescale time $\tilde{t} = c_1 t$ and define effective parameters,

$$\beta_1 = \alpha_2 k_1, \quad \beta_2 = \frac{\alpha_3\alpha_4 k_2}{\alpha_5}, \quad \beta_3 = \frac{k_3}{\alpha_5}, \quad \beta_4 = \frac{k_4}{\alpha_5}, \quad \beta_5 = \frac{k_5}{\alpha_5}, \quad \beta_6 = \alpha_1\alpha_5,$$

$$\nu_2 = \frac{a_2}{c_2},\ \nu_3 = \frac{a_3}{c_3},\ \nu_4 = \frac{a_4}{c_4},\ \nu_5 = \frac{a_5}{c_5}.$$

The rescaled system takes the form

$$\boldsymbol{T}\frac{\partial \boldsymbol{w}}{\partial t} = \boldsymbol{D}\frac{\partial^2 \boldsymbol{w}}{\partial x^2} + \boldsymbol{F}(\boldsymbol{w}, \boldsymbol{\beta}), \tag{S2}$$

where

$$\boldsymbol{w} = \left([W_l](x,t), [A](x,t), [W_d](x,t), [C](x,t), [S](x,t)\right), \tag{S3}$$

$$\boldsymbol{T} = diag(1, \tau_2, \dots, \tau_5), \tau_i = \frac{c_1}{c_i}, i = 2, \dots, 5, \quad \boldsymbol{D} = diag(0, \nu_2, \dots, \nu_5), \tag{S4}$$

$$\boldsymbol{F}(\boldsymbol{w}, \boldsymbol{\beta}) = \begin{pmatrix} \dfrac{\beta_6[S]}{(1+[A])(1+[C])(1+\beta_3[W_l])} - [W_l] \\ \dfrac{\beta_1}{(1+\beta_4[W_l])} - [A] \\ \beta_2[W_l][S] - [W_d] \\ \dfrac{[W_d]}{(1+\beta_5[W_l])} - [C] \\ [W_l] - [S] \end{pmatrix}, \tag{S5}$$

with $\boldsymbol{\beta} = (\beta_1, \dots, \beta_6)$ being the obtained above non-dimensionalised effective model parameters.

#### Spatially homogeneous steady states and their stability

Solving the algebraic equations for spatially uniform steady states, we obtain

$$\beta_6[S] = [W_l](1+[A])(1+[C])(1+\beta_3[W_l]), \tag{S6}$$

with

$$[A] = \frac{\beta_1}{1+\beta_4[W_l]},\ [W_d] = \beta_2[W_l]^2,\ [C] = \frac{\beta_2[W_l]^2}{1+\beta_5[W_l]},\ [S] = [W_l]. \tag{S7}$$

Substituting (S7) to (S6) and denoting $w := [W_l]$ results in

$$\beta_6 w = w\left(1 + \frac{\beta_1}{1+\beta_4 w}\right)\left(1 + \frac{\beta_2 w^2}{1+\beta_5 w}\right)(1+\beta_3 w). \tag{S8}$$

Eq. (S8) has a trivial solution $w = 0$, which corresponds to the semi-trivial steady state $\boldsymbol{w}_0$ with components

$$[W_l] = 0,\ [A] = \beta_1,\ [W_d] = 0,\ [C] = 0,\ [S] = 0. \tag{S9}$$

Positive homogeneous steady states satisfies

$$\beta_6(1+\beta_4 w)(1+\beta_5 w) = (1+\beta_1+\beta_4 w)(1+\beta_5 w+\beta_2 w^2)(1+\beta_3 w), \tag{S10}$$

and can be found as the intersection points of $f_1(w)$ and $f_2(w)$, where

$$f_1(w) = \beta_6(1+\beta_4 w)(1+\beta_5 w), \tag{S11}$$

and

$$f_2(w) = (1+\beta_1+\beta_4 w)(1+\beta_5 w+\beta_2 w^2)(1+\beta_3 w). \tag{S12}$$

Depending on model parameters, there can be a different number of intersections of these functions (Figure 1). It holds

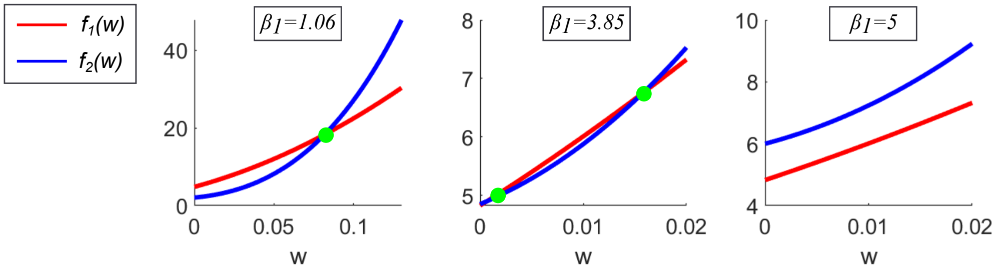


**Fig. S 1.** Intersections of functions $f_1(w)$ and $f_2(w)$ for different values of model parameters $\beta_i$, $i = 1, \dots, 6$. The value of parameter $\beta_1$ is given in each sub-figure, while all other parameters are fixed to the values given in Table 1.

**Lemma S1.** *The reduced model* (S2) *has no more than three non-trivial spatially homogeneous steady states.*

*Proof.* $f(w) = f_2(w) - f_1(w) = 0$ is a polynomial with $Deg(f(w)) = 4$. Thus, depending on model parameters, it can have no more than four roots. Furthermore, taking into account that coefficients at the two highest variable exponents are positive and applying Descartes' rule of signs, we conclude that the maximal number of coefficient sign changes and positive real roots is equal to three. □

Linearisation of (S2) at $w_0$ results in

$$\boldsymbol{T}\frac{\partial \boldsymbol{w}}{\partial t} = \boldsymbol{D}\frac{\partial^2 \boldsymbol{w}}{\partial x^2} + \boldsymbol{N_1}(\boldsymbol{\beta})\boldsymbol{w}, \tag{S13}$$

where

$$\boldsymbol{N_1}(\boldsymbol{\beta}) = \frac{\partial \boldsymbol{N_1}(\boldsymbol{w}, \boldsymbol{\beta})_i}{\partial \boldsymbol{w}_j}|_{\boldsymbol{w}=\boldsymbol{w}_0} = \begin{pmatrix} -1 & 0 & 0 & 0 & \frac{\beta_6}{1+\beta_1} \\ -\beta_1\beta_4 & -1 & 0 & 0 & 0 \\ 0 & 0 & -1 & 0 & 0 \\ 0 & 0 & 1 & -1 & 0 \\ 1 & 0 & 0 & 0 & -1 \end{pmatrix}. \tag{S14}$$

We define a linear operator

$$L_N := \{\boldsymbol{D}\frac{\partial^2}{\partial x^2} + \boldsymbol{N_1} \text{ with homogeneous Neumann boundary conditions}\} \tag{S15}$$

Applying the spectral decomposition of the Laplace operator with homogeneous Neumann boundary conditions in $L^2(\Omega)$, we can characterise the spectrum of $L_N$ by the spectrum of

$$\tilde{\boldsymbol{N_1}} := -\boldsymbol{D}\lambda_k + \boldsymbol{N_1},$$

where $\lambda_k = (\pi k)^2$, $k = 0, 1, \dots$ , and $\psi_0(x) = 1$, $\psi_k(x) = \sqrt{2}\cos(\pi k x), k = 1, 2, \dots$, denote the eigenvalues and eigenfunctions of the Laplace operator $-\frac{\partial^2}{\partial x^2}$, defined on the interval $(0, 1)$ with homogeneous Neumann boundary conditions,

$$-\frac{d^2}{dx^2}\psi_k(x) = \lambda_k\psi_k(x), \quad \frac{d}{dx}\psi_k(0) = \frac{d}{dx}\psi_k(1) = 0, \quad k = 0, 1, \dots. \tag{S16}$$

This reduces the linearised stability problem to the analysis of the characteristic polynomials

$$\varphi_k(\sigma) = det(\boldsymbol{T}^{-1}(\boldsymbol{N_1}(\boldsymbol{\beta}) - \lambda_k\boldsymbol{D}) - \sigma_k I) = 0, \quad k = 0, 1, \dots, \tag{S17}$$

which can be rewritten as

$$\varphi_k(\sigma) = -(\sigma+\frac{1+\nu_2\lambda_k}{\tau_2})(\sigma+\frac{1+\nu_3\lambda_k}{\tau_3})(\sigma+\frac{1+\nu_4\lambda_k}{\tau_4})(\sigma^2+(1+\frac{1+\nu_5\lambda_k}{\tau_5})\sigma+\frac{1+\nu_5\lambda_k}{\tau_5}-\frac{\beta_6}{\tau_5(1+\beta_1)}). \tag{S18}$$

It yields the stability conditions

$$1+\nu_5\lambda_k - \frac{\beta_6}{1+\beta_1} > 0, \quad k = 0, 1, \ldots,$$

which for $k = 0$ read,

$$1 - \frac{\beta_6}{1+\beta_1} > 0 \Rightarrow \beta_6 < 1 + \beta_1.$$

Consequently, we conclude that the semi-trivial steady state (S9) is always locally asymptotically stable or unstable, i.e. cannot exhibit the Turing instability.

**Lemma S2.** *If trivial steady state* (S9) *is linearly unstable, then there exists at least one non-trivial spatially homogeneous steady state.*

*Proof.* The sufficient condition for linear instability reads

$$\beta_6 > 1 + \beta_1.$$

However, in this case

$$f_1(0) = \beta_6, \quad f_2(0) = 1 + \beta_1,$$

and we have $f_1(0) > f_2(0)$. As $Deg(f_1(w)) = 2$, $Deg(f_2(w)) = 4$, and both polynomials have positive coefficients at leading variable exponents, functions $f_1(w)$ and $f_2(w)$ must have at least one intersection for $w > 0$. □

**Lemma S3.** *Let $\beta_4 \leq \beta_5$, $\beta_3 < \beta_5$, and $\beta_6 > 1 + \beta_1$. Then, model* (S1) *has exactly one non-trivial spatially homogeneous steady state.*

*Proof.* Let $w^1 = -\frac{1}{\beta_4}$ and $w^2 = -\frac{1+\beta_1}{\beta_4}$, so that $w^2 < w^1$. Under the assumptions on the limiting constants, we have $f_1(w^1) = 0$ and $f_2(w^1) > 0$, while $f_1(w^2) > 0$ and $f_2(w^2) = 0$. Since both functions are continuous, there must exist at least one point of intersection of $f_1(w)$ and $f_2(w)$ in the interval $[w^2, w^1]$.

Furthermore, since $\deg(f_1(w)) = 2$, $\deg(f_2(w)) = 4$ and both polynomials have positive leading coefficients, additional intersection must occur for $w < w^2$. Additionally, under the assumptions on the model parameters, we have $f_2(0) < f_1(0)$, indicating another point of intersection for negative values of $w$. Therefore, there can be at most one intersection of $f_1(w)$ and $f_2(w)$ for positive values of $w$. From Lemma S2, it follows that such an intersection indeed exists. □

**Lemma S4.** *Let*

$$\begin{aligned} &\beta_1 + 1 > \beta_6; \\ &\beta_4 + (\beta_3 + \beta_5)(\beta_1 + 1) > \beta_6(\beta_4 + \beta_5); \\ &(\beta_1 + 1)(\beta_2 + \beta_3\beta_5) + \beta_4(\beta_3 + \beta_5) > \beta_4\beta_5\beta_6. \end{aligned} \tag{S19}$$

*Then, there are no non-trivial steady states of model* (S2).

*Proof.* Let us consider the function used to identify homogeneous steady states: $f(w) = f_2(w) - f_1(w)$. The conditions specified in (S19) ensure that all coefficients of this polynomial are positive. As a consequence, the polynomial $f(w)$ has no positive real roots. Hence, the system admits only the trivial steady state. Furthermore, this steady state is linearly stable, as guaranteed by the first condition in (S19). It follows that all solutions of the model asymptotically approach the trivial steady state. □

| Parameter name | Parameter value | Parameter name | Parameter value |
|---|---|---|---|
| $\nu_1$ | 0 | $\beta_1$ | 1.06 |
| $\nu_2$ | $3.81 \times 10^{-5}$ | $\beta_2$ | 540.4 |
| $\nu_3$ | $4.43 \times 10^{-1}$ | $\beta_3$ | 1.15 |
| $\nu_4$ | $6.07 \times 10^{-8}$ | $\beta_4$ | 11.59 |
| $\nu_5$ | $4.78 \times 10^{-4}$ | $\beta_5$ | 11.59 |
| $\tau_i,\ i = 1, \ldots, 5$ | 1 | $\beta_6$ | 4.82 |

**Tab. S 1.** Values of model parameters, obtained by fitting the one-dimensional model (S2) to the 3D pattern data obtained from numerical integration of the 3D model.

**Corollary S1.** *Let us assume that the following conditions are satisfied:*

$$\begin{aligned} &\beta_1 + 1 < \beta_6, \\ &(\beta_1 + 1)\,(\beta_2 + \beta_3\,\beta_5) + \beta_4\,(\beta_3 + \beta_5) > \beta_4\,\beta_5\,\beta_6. \end{aligned} \tag{S20}$$

*Under these conditions, there exists exactly one non-trivial steady state. In particular, the system does not admit bistability.*

*Proof.* The conditions in (S20) ensure that the function $f(w)$ is strictly convex and satisfies $f(0) < 0$. Consequently, $f(w)$ has exactly one positive real root. □

**Comparing one-dimensional model with three-dimensional simulations**

Before conducting a numerical analysis of pattern formation, it is necessary to determine parameter values for the one-dimensional model (S2) that yield patterns qualitatively similar to those observed in 3D model simulations. The model fitting procedure is as follows. First, the data obtained from the 3D model simulations are pre-processed by averaging the concentration values in the direction perpendicular to the body axis and scaling the resulting one-dimensional profiles to the unit interval using min-max normalisation. This produces data that is suitable for fitting the one-dimensional model; here, our primary interest is in matching the shapes of the gradient profiles rather than the precise concentration values.

We define the residual between the output of (S2) and the pattern data in the least-squares sense,

$$S(\boldsymbol{\nu}, \boldsymbol{\beta}) = \sum_{i=1}^{M_{\dim}} \left\| \bar{\boldsymbol{w}}_i(\boldsymbol{\nu}, \boldsymbol{\beta}) - \bar{\boldsymbol{w}}_i^0 \right\|^2, \tag{S21}$$

where $\bar{\boldsymbol{w}}_i^0$ denotes the pre-processed pattern data from the 3D model evaluated at the grid points $x_i$, for $i = 1, \ldots, M_{\dim}$, and $\bar{\boldsymbol{w}}_i(\boldsymbol{\nu}, \boldsymbol{\beta})$ is the numerical approximation of the stationary solution of the spatially discretised one-dimensional model with the corresponding parameter values. The initial conditions for the numerical simulation are defined by applying small perturbations to the homogeneous steady state for all model components except for the source density, for which a gradient is prescribed

$$[SD_0](x) = [SD_h] + 4\,\exp(x),$$

where $[SD_h]$ is the corresponding component of the homogeneous steady state. This enforces convergence of the numerical solution to the desired pattern, thereby reducing the variability of the residual function $S(\boldsymbol{\nu}, \boldsymbol{\beta})$ and simplifying the optimisation process. Subsequently, we minimise the residual function (S21) to obtain optimal values for the diffusion coefficients $\nu_i$ and reaction rates $\beta_i$, for $i = 1, \ldots, 5$, corresponding to a local minimum of the residual. In this step, we set $\tau_i = 1$, for $i = 1, \ldots, 5$, to reduce the dimensionality of the optimisation problem. The estimated model parameters are listed in Table 1, and the resulting concentration profiles are illustrated in Figure 2.

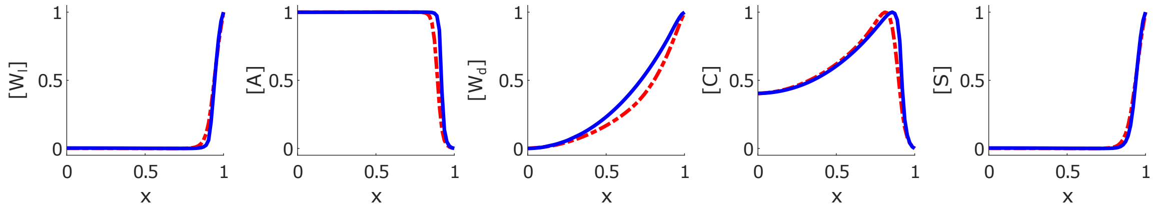


**Fig. S 2.** Steady-state pattern obtained from the numerical simulation of the one-dimensional model (S2) with the parameter values from Table 1 (blue) and concentration profiles from the 3D simulations, averaged in the direction perpendicular to the body axis (red). Both concentration profiles are scaled to the interval $[0, 1]$ with the min-max normalisation procedure.

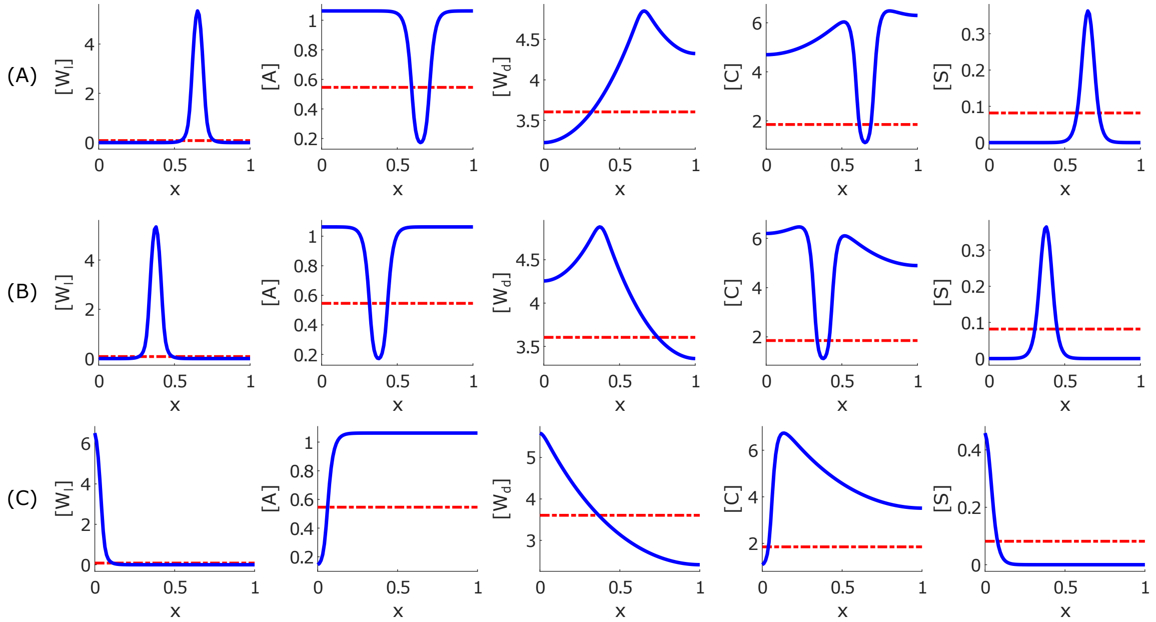


**Fig. S 3.** Three examples of pattern formation in the one-dimensional model (A-C). Parameter values used in the simulations are given in Table 1. Initial data are taken as small perturbations of the homogeneous steady state. Blue colour shows the final concentration profiles, while red colour denotes the initial concentration values.

The obtained model parameters satisfy the conditions of Lemma S3. Consequently, the trivial steady state (S9) is linearly unstable, and there exists a unique non-trivial spatially homogeneous steady state, denoted by $\boldsymbol{w}_1$, which can be computed numerically as

$$\boldsymbol{w}_1 \approx (0.08,\ 0.54,\ 3.60,\ 1.85,\ 0.08).$$

Numerical analysis reveals that $\boldsymbol{w}_1$ exhibits a Turing instability. In all our numerical experiments, regardless of the initial data chosen, the numerical solution consistently converged to spatially inhomogeneous patterns. The resulting concentration profiles are shown in Figure 2. However, the shape of the resulting patterns may depend on the initial data, as illustrated in Figure 3.

##### Semi-analytical approach to the analysis of branching patterns

To decipher the nature of the patterns observed in simulations, we analyse the stability of branching stationary solutions near the Turing bifurcation point. As the complexity of the model equations precludes a fully rigorous analytical treatment, we adopt a hybrid approach that combines analytical techniques with numerical methods. In particular, we compute numerical approximations for quantities that cannot be determined analytically, such as the coordinates of steady states and the eigenvalues of the linearised system.

We apply the classical Lyapunov-Schmidt method [1, 2] to investigate the branching behaviour of stationary solutions of the one-dimensional model (S2). To streamline the analysis, we set $\boldsymbol{T} = \boldsymbol{I}$. We assume that all parameters are fixed except for the largest diffusion coefficient, $\nu_3 := \nu$, which we treat as a bifurcation parameter. Expanding the nonlinearity $\boldsymbol{N}(\boldsymbol{w}, \boldsymbol{\beta})$ in a Taylor series about a spatially homogeneous steady state $\boldsymbol{w}_1$, we obtain

$$\frac{\partial \boldsymbol{u}}{\partial t} = \boldsymbol{D}(\nu)\frac{\partial^2 \boldsymbol{u}}{\partial x^2} + \boldsymbol{N}_1(\boldsymbol{u}) + \boldsymbol{N}_2(\boldsymbol{u}, \boldsymbol{u}) + \boldsymbol{N}_3(\boldsymbol{u}, \boldsymbol{u}, \boldsymbol{u}) + \dots, \quad \boldsymbol{w} = \boldsymbol{w}_1 + \boldsymbol{u}, \tag{S22}$$

where the linear operator $\boldsymbol{N}_1$ is defined by the Jacobian matrix (S14), and the higher-order terms $\boldsymbol{N}_2$ and $\boldsymbol{N}_3$ are given by

$$(\boldsymbol{N}_2)_j(\boldsymbol{u}, \boldsymbol{u}) = \frac{1}{2} \sum_{i_1, i_2 = 1}^{5} u_{i_1} u_{i_2} \left. \frac{\partial^2 \boldsymbol{N}_j}{\partial u_{i_1} \partial u_{i_2}} \right|_{\boldsymbol{u}=0}, \quad j = 1, \dots, 5,$$

$$(\boldsymbol{N}_3)_j(\boldsymbol{u}, \boldsymbol{u}, \boldsymbol{u}) = \frac{1}{6} \sum_{i_1, i_2, i_3 = 1}^{5} u_{i_1} u_{i_2} u_{i_3} \left. \frac{\partial^3 \boldsymbol{N}_j}{\partial u_{i_1} \partial u_{i_2} \partial u_{i_3}} \right|_{\boldsymbol{u}=0}, \quad j = 1, \dots, 5.$$

Next, we consider a linear spectral problem for the operator $L_N(\nu)$, defined in (S15), on $L^2(\Omega)$,

$$L_N(\nu)\boldsymbol{\phi} = \sigma\boldsymbol{\phi}, \ \boldsymbol{\phi} \neq 0, \tag{S23}$$

and seek an eigenfunction $\boldsymbol{\phi}$ in the form of a series expansion with vector coefficients,

$$\boldsymbol{\phi} = \sum_{k=0}^{+\infty} \boldsymbol{C}_k \psi_k(x), \quad \boldsymbol{C}_k \in \mathbb{R}^5. \tag{S24}$$

By substituting (S24) into (S23) and matching coefficients of identical eigenfunctions, we derive a sequence of linear systems,

$$\boldsymbol{N}_1 - \boldsymbol{D}(\nu)\lambda_k \boldsymbol{I} = 0, \quad k = 0, 1, \dots, \tag{S25}$$

from which the eigenvalues $\sigma_j^k(\nu)$, for $j = 1, \dots, 5$, of the operator $L_N(\nu)$ can be determined.

The critical value $\nu_{\text{cr}}$ of the control parameter $\nu$ is defined as the value at which the spectrum of the linear operator $L_N(\nu_{\text{cr}})$ lies entirely within the closed left half-plane of the complex plane, with at least one eigenvalue $\sigma$ located on the imaginary axis and satisfying

$$\left. \frac{d\,\text{Re}(\sigma)}{d\nu} \right|_{\nu=\nu_{\text{cr}}} \neq 0.$$

We recall that the spatially homogeneous steady state $\boldsymbol{w}_1$ is said to be *Turing unstable* if the following conditions are satisfied:

1. All eigenvalues $\sigma_j^0$, for $j = 1, \dots, 5$, of the linearised system *without* diffusion lie strictly in the left half-plane of the complex plane.

2. There exists at least one eigenvalue $\sigma_{j_0}^{k_0}$, with $k_0 > 0$, of the linearised system *with* diffusion that lies in the right half-plane of the complex plane.

If the only eigenvalue on the imaginary axis at $\nu = \nu_{\text{cr}}$ is $\sigma = 0$, the system undergoes a *monotonic* (or *stationary*) instability. In contrast, if a pair of purely imaginary eigenvalues $\sigma = \pm i\omega_0$ exists at $\nu = \nu_{\text{cr}}$, the system exhibits an *oscillatory* instability, commonly referred to as a *Hopf bifurcation.*

Assuming that $\nu_{\text{cr}}$ is the critical value of the control parameter $\nu$ such that there exists a single simple eigenvalue $\sigma_{j_0}^{k_0}(\nu_{\text{cr}}) = 0$, while all other eigenvalues lie strictly in the left half-plane of the complex plane, we construct an asymptotic approximation of the branching stationary solutions using the Lyapunov–Schmidt reduction method. As a first step, we seek the eigenfunctions of the linearised spectral problem and its adjoint:

$$L_N(\nu_{\text{cr}})\boldsymbol{\varphi} = 0, \quad L_N^*(\nu_{\text{cr}})\boldsymbol{\Phi} = 0, \quad \langle \boldsymbol{\varphi}, \boldsymbol{\Phi} \rangle = 1,$$

where $\boldsymbol{\varphi}$ and $\boldsymbol{\Phi}$ are the eigenfunctions of the operator $L_N(\nu_{\text{cr}})$ and its adjoint $L_N^*(\nu_{\text{cr}})$, respectively. The inner product is normalised such that $\langle \boldsymbol{\varphi}, \boldsymbol{\Phi} \rangle = 1$. Due to the simplicity of $\sigma_{j_0}^{k_0}(\nu_{\text{cr}})$, these functions can be expressed as

$$\boldsymbol{\varphi} = \boldsymbol{b}\,\psi_{k_0}(x), \quad \boldsymbol{\Phi} = \boldsymbol{c}\,\psi_{k_0}(x). \tag{S26}$$

Next, we define in (S22) the small parameter $\varepsilon^2 = \nu - \nu_{\text{cr}}$, set $\boldsymbol{u}_t = 0$, and obtain the stationary equation,

$$0 = L_N(\nu_{\text{cr}})\boldsymbol{u} + \varepsilon^2 \boldsymbol{B}\boldsymbol{u} + \boldsymbol{N}_2(\boldsymbol{u}, \boldsymbol{u}) + \boldsymbol{N}_3(\boldsymbol{u}, \boldsymbol{u}, \boldsymbol{u}) + \ldots, \tag{S27}$$

where $\boldsymbol{B}\boldsymbol{u} = \boldsymbol{D}_1 \dfrac{d^2\boldsymbol{u}}{dx^2}$ and $\boldsymbol{D}_1 = \text{diag}(0, 0, 1, 0, 0)$.

Branching stationary solutions for $\varepsilon > 0$ are constructed in the form of a power series:

$$\boldsymbol{u} = \sum_{k=1}^{\infty} \varepsilon^k \boldsymbol{u}_k. \tag{S28}$$

By substituting (S28) into (S27) and equating the coefficients of like powers of $\varepsilon$, we obtain the following chain of equations:

$$\varepsilon^1 : \quad -L_N(\nu_{\text{cr}})\boldsymbol{u}_1 = 0; \tag{S29}$$

$$\varepsilon^2 : \quad -L_N(\nu_{\text{cr}})\boldsymbol{u}_2 = \boldsymbol{N}_2(\boldsymbol{u}_1, \boldsymbol{u}_1) \equiv \boldsymbol{f}_2; \tag{S30}$$

$$\varepsilon^3 : \quad -L_N(\nu_{\text{cr}})\boldsymbol{u}_3 = \boldsymbol{B}\boldsymbol{u}_1 + 2\boldsymbol{N}_2(\boldsymbol{u}_1, \boldsymbol{u}_2) + \boldsymbol{N}_3(\boldsymbol{u}_1, \boldsymbol{u}_1, \boldsymbol{u}_1) \equiv \boldsymbol{f}_3. \tag{S31}$$

The solution to equation (S29) is given by the expression:

$$\boldsymbol{u}_1 = \beta_1 \boldsymbol{\varphi},$$

where the constant $\beta_1$ will be determined at a later stage of the method. The solvability condition for the inhomogeneous equation (S30) is

$$(\boldsymbol{f}_2, \boldsymbol{\Phi}) = \int_0^1 \left( f_2^1 \Phi_1 + \ldots + f_2^5 \Phi_5 \right) dx = 0,$$

which is automatically satisfied due to the structure of $\boldsymbol{f}_2$. The solution of equation (S30) can then be written as

$$\boldsymbol{u}_2 = \beta_2 \boldsymbol{\varphi} + \boldsymbol{u}_{20}, \quad \boldsymbol{u}_{20} = \beta_1^2 \left( \boldsymbol{v}_2^0 + \boldsymbol{v}_2^{2k_0} \psi_{2k_0}(x) \right),$$

where the vectors $\boldsymbol{v}_2^0$ and $\boldsymbol{v}_2^{2k_0}$ are determined as solutions of the following linear systems:

$$N_1 \boldsymbol{v}_2^0 = -\frac{1}{2} N_2(\boldsymbol{b}, \boldsymbol{b}), \quad (N_1 - \lambda_{2k_0} \boldsymbol{D}(\nu_{\text{cr}}))\, \boldsymbol{v}_2^{2k_0} = -\frac{1}{2} N_2(\boldsymbol{b}, \boldsymbol{b}).$$

The solvability condition for equation (S31) takes the form:

$$(\boldsymbol{f}_3, \boldsymbol{\Phi}) = \beta_1(B\boldsymbol{\varphi}, \boldsymbol{\Phi}) + 2\beta_1\beta_2(N_2(\boldsymbol{\varphi}, \boldsymbol{\varphi}), \boldsymbol{\Phi}) + 2\beta_1^3 \left(N_2\left(\boldsymbol{\varphi}, \boldsymbol{v}_2^0 + \boldsymbol{v}_2^{2k_0}\psi_{2k_0}(x)\right), \boldsymbol{\Phi}\right) + \beta_1^3(N_3(\boldsymbol{\varphi}, \boldsymbol{\varphi}, \boldsymbol{\varphi}), \boldsymbol{\Phi}) = 0.$$

From this, we derive the expression for $\beta_1^2$:

$$\beta_1^2 = -\frac{(B\boldsymbol{\varphi}, \boldsymbol{\Phi})}{(N_3(\boldsymbol{\varphi}, \boldsymbol{\varphi}, \boldsymbol{\varphi}), \boldsymbol{\Phi}) + 2(N_2(\boldsymbol{\varphi}, \boldsymbol{v}_2^0 + \boldsymbol{v}_2^{2k_0}\psi_{2k_0}(x)), \boldsymbol{\Phi})}, \tag{S32}$$

where

$$(N_3(\boldsymbol{\varphi}, \boldsymbol{\varphi}, \boldsymbol{\varphi}), \boldsymbol{\Phi}) = \frac{3}{8}(N_3(\boldsymbol{b}, \boldsymbol{b}, \boldsymbol{b}), \boldsymbol{c}),$$

$$(N_2(\boldsymbol{\varphi}, \boldsymbol{v}_2^0 + \boldsymbol{v}_2^{2k_0}\psi_{2k_0}(x)), \boldsymbol{\Phi}) = \frac{1}{2}(N_2(\boldsymbol{b}, \boldsymbol{v}_2^0), \boldsymbol{c}) + \frac{1}{4}(N_2(\boldsymbol{b}, \boldsymbol{v}_2^{2k_0}), \boldsymbol{c}),$$

$$(B\boldsymbol{\varphi}, \boldsymbol{\Phi}) = \frac{1}{2}(-\lambda_{k_0} D_1 \boldsymbol{b}, \boldsymbol{c}).$$

Here, the sign of $\beta_1^2$ determines the linear stability of the branching solutions. If $\beta_1^2$ is positive, then the pair of branching stationary solutions

$$\boldsymbol{w}_{1,2} = \boldsymbol{w}_1 \pm \varepsilon\beta_1\boldsymbol{\varphi} + O(\varepsilon^2)$$

is stable for small $\varepsilon = \sqrt{\nu - \nu_{\mathrm{cr}}}$.

To apply the method, we fix the model parameters to the values given in Table 1. The steady state $\boldsymbol{w}_1$ is found using Newton's method:

$$\boldsymbol{w}_1 \approx (0.0816, 0.5458, 3.6049, 1.8513, 0.0816).$$

The critical value of the control parameter $\nu_{\mathrm{cr}}$ is estimated by considering the eigenvalue problem (S23) and truncating the series expansion (S24) with a sufficiently large cut-off value $N = 100$:

$$\boldsymbol{u} \approx \sum_{k=0}^{N} \boldsymbol{C}_k \phi_k(x). \tag{S33}$$

Substituting (S33) into (S23), we obtain a finite number of linear systems:

$$\boldsymbol{N}_1 - \boldsymbol{D}(\nu)\lambda_k \boldsymbol{I} = 0, \quad k = 0, 1, 2, \ldots, N,$$

from which we numerically determine the eigenvalues $\sigma_j^k(\nu)$. We introduce the residual function

$$f(\nu) = \max\left|\sigma_j^k(\nu)\right|, \quad k = 0, \ldots, N,\ j = 1, \ldots, 5,$$

and minimize it with respect to $\nu$ using standard nonlinear optimization methods. The result is

$$\nu_{\mathrm{cr}} \approx 0.0094.$$

The first eigenvalues of the operator $L_N(\nu_{\mathrm{cr}})$ are shown in Figure 4, part A. All plotted values are negative, except for one eigenvalue $\sigma_j^{k_0} \approx 0$, where $k_0 = 8$. Next, we estimate the vectors $\boldsymbol{b}$ and $\boldsymbol{c}$ in the expressions for eigenfunctions (S26),

$$b \approx (-0.2280, 0.7238, -2.5573, 1.2007, -0.1751), \quad c \approx (-2.4022, 0.1239, 0.0050, 0.0688, -1.6726).$$

Next, we evaluate the expression (S32) and obtain $\beta_1^2 \approx 44.6539 > 0$, meaning that branching stationary solutions are stable (see Figure 4, part B). This corresponds to the scenario of Turing pattern formation.

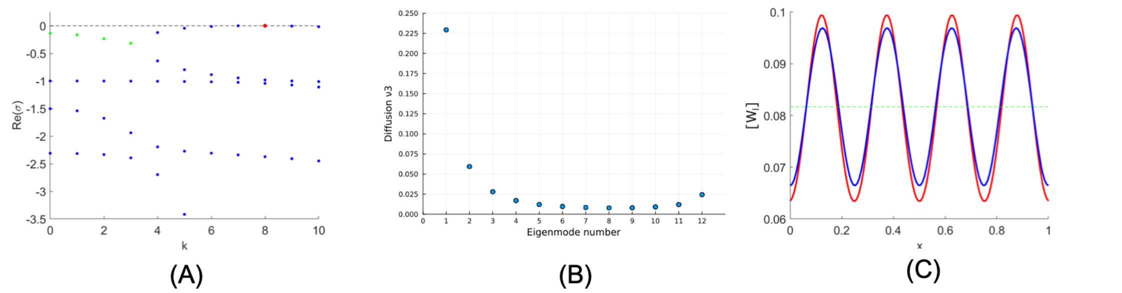


**Fig. S 4.** The results of a semi-analytical bifurcation analysis of the one-dimensional system. (A): The real parts of the eigenvalues of the linear operator $L_N(\nu_{\rm cr})$. Real eigenvalues are shown in blue, while the real parts of complex eigenvalues are depicted in green. The critical eigenvalue is highlighted in red. (B) Unstable modes in dependence of the diffusion coefficient of $[W_d]$. (C): The asymptotics of the secondary stationary solution (first component), evaluated at $\varepsilon = 0.01$ (blue), compared to the numerical solution of the system computed with the same parameter values (red). The green dashed line represents the respective component of the spatially homogeneous steady state $\boldsymbol{w}_1$. The initial conditions for the simulation are small random perturbations of the stationary state.

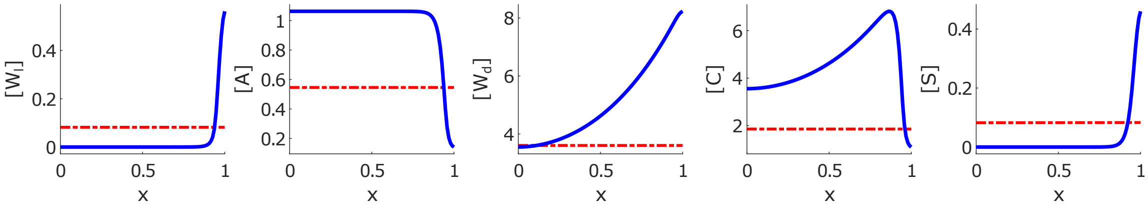


**Fig. S 5.** Example pattern, obtained by numerically integrating system (S34)-(S38). Initial data are taken as small perturbations of the homogeneous steady state $\boldsymbol{w_1}$. The blue colour shows the final concentration profiles, while the red colour denotes the initial concentration values.

#### Model variations

To examine the robustness of the one-dimensional model (S1) to changes in reaction terms, we consider two variations of the model. The first version is obtained by multiplying the reaction terms in (S1) by the respective degradation terms, giving the following system of equations,

$$\partial_t[W_l] = \quad b_1[S] - c_1[W_l](1 + k_1[A])(1 + k_2[C])(1 + k_3[W_l]) \tag{S34}$$

$$\partial_t\,[A] = \quad a_2\frac{\partial}{\partial x^2}[A] + b_2 - c_2[A](1 + k_4[W_l]) \tag{S35}$$

$$\partial_t[W_d] = \quad a_3\frac{\partial}{\partial x^2}[W_d] + b_3[W_l][S] - c_3[W_d] \tag{S36}$$

$$\partial_t[C] = \quad a_4\frac{\partial}{\partial x^2}[C] + b_4[W_d] - c_4[C](1 + k_5[W_l]) \tag{S37}$$

$$\partial_t[S] = \quad a_5\frac{\partial}{\partial x^2}[S] + b_5[W_l] - c_5[S]. \tag{S38}$$

It can be interpreted as accounting for regulatory feedback in the decay terms.

System (S34)-(S38) has obviously the same spatially homogeneous steady states as the original system (S1). Next, we simulate this system for the parameter values shown in Table 1. An example pattern is shown in Figure 5.

Another modification was obtained by replacing some reaction terms in system (S1) with higher

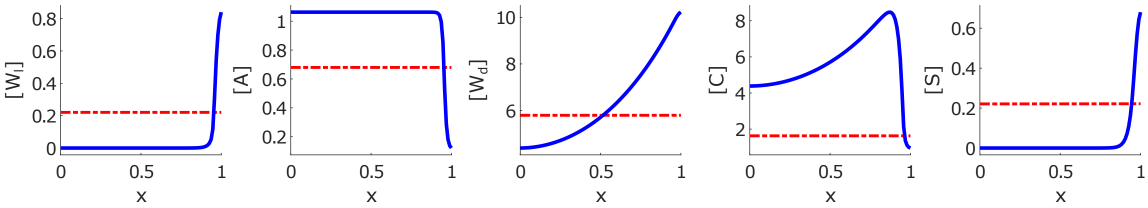


**Fig. S 6.** Example pattern, obtained by numerically integrating system (S39)-(S43). Initial data are taken as small perturbations of the homogeneous steady state $\boldsymbol{w_1}$. The blue colour shows the final concentration profiles, while the red colour denotes the initial concentration values.

power terms. We arrive at the following system of equations

$$\partial_t[W_l] = \frac{b_1[S]}{(1+k_1[A]^2)(1+k_2[C])(1+k_3[W_l])} - c_1[W_l] \tag{S39}$$

$$\partial_t\,[A] = a_2\frac{\partial}{\partial x^2}[A] + \frac{b_2}{(1+k_4[W_l]^2)} - c_2[A] \tag{S40}$$

$$\partial_t[W_d] = a_3\frac{\partial}{\partial x^2}[W_d] + b_3[W_l][S]^2 - c_3[W_d] \tag{S41}$$

$$\partial_t[C] = a_4\frac{\partial}{\partial x^2}[C] + \frac{b_4[W_d]}{(1+k_5[W_l])} - c_4[C] \tag{S42}$$

$$\partial_t[S] = a_5\frac{\partial}{\partial x^2}[S] + b_5[W_l] - c_5[S]. \tag{S43}$$

We simulate the model (S39)-(S43) using parameters from Table 1. The resulting pattern is shown in Figure 6.

#### Mathematical framework for modelling pseudo 3D geometry

The computational work presented in this paper is based on a mechano-chemical modelling framework [3–5], which couples the Mutual Inhibition Model (MI) of Wnt-Dkk interactions, described by the reaction-diffusion equations in Eq. (1)-(5) (main manuscript), to a continuous tissue evolution model.

The tissue, forming a hollow ellipsoid in the shape of a *Hydra* cell bilayer, is approximated at each time $t$ by a closed 2D surface $\Gamma(t)$ embedded in 3D space. Detailed modelling of the cell layers is neglected. The evolution of $\Gamma(t)$ is represented by a diffeomorphic, time-dependent map $\vec{X}$, parameterised over the unit sphere $S^2 \subset \mathbb{R}^3$, so that $\Gamma(t)$ is the image of $\vec{X}(\cdot,t)$, with $\vec{X}(\vec{s},t) : S^2 \times [0,T] \to \mathbb{R}^3$ for $T \in \mathbb{R}_{>0}$. The chemical processes on the evolving tissue surface are modelled by identifying material points $\vec{X}(\vec{s},t)$ on $\Gamma(t)$ with $\vec{s} \in S^2$. The smooth and bijective nature of $\vec{X}$ ensures this correspondence. Functions $\phi_a : S^2 \times [0,T] \to \mathbb{R}_{\geq 0}$, given by $\phi_a(\vec{s},t) = \Phi_a(\vec{X}(\vec{s},t))$, describe the local concentrations of gene products, where $\Phi_a$ represents the concentrations $([W_l],[A],[W_d],[C],[S])$ in the MI model.
Assuming purely elastic tissue, the elastic deformations are modelled using Helfrich's free energy [6], defined as:

$$\mathcal{F}_{bend} = \int_\Gamma \kappa\big(H - H_0([W_l],[T_a],[F_a])\big)^2\, d\vec{S},$$

where $H$ is the mean curvature, $\kappa$ is the bending rigidity, and $H_0$ represents the spontaneous curvature [3,4]. $H_0$ is assumed to depend on local morphogen concentrations, and we take
$H_0([W_l],[T_a],[F_a]) = 1.5[W_l] + 3.0[F_a] + 10.0[T_a]$,
based on local tissue evaginations observed during head, foot, and tentacle formation [7–9]. The

evolution of the tissue surface is computed from the $L^2$-gradient flow of the total energy, including a local Lagrange multiplier. Minimization of the free energy results in a 4th order PDE model for the deforming tissue surface. This model is coupled to the reaction-diffusion system in Eq. (1)-(5) (main manuscript), with the diffusion process represented by the surface Laplace-Beltrami operator $\Delta^\Gamma$. For further details, see Ref. [3].

**Extended model for tentacle formation**

To complement the qualitative description in the main text, we provide here the explicit mathematical formulation of the tentacle subsystem. The tentacle module is phenomenological and represented by an activator–inhibitor system receiving positive input from the source density ($[S]$) and negative input from local Wnt activity ($[W_l]$). This structure captures the experimentally observed formation of tentacle primordia in regions of intermediate head-forming competence, where $[S]$ is high but $\beta$-catenin/Wnt locally suppresses/shifts the tentcale system [10–13]. Including the tentacle subsystem does not affect Wnt–Dkk pattern formation but provides additional data for model verification, such as the response to ALP treatment.

$$d_t[T_a] = a_6\Delta^\Gamma[T_a] + \frac{b_6[S](g+[T_a]^2)}{[T_i](e+d[T_a]^2)(e+h[W_l])} - c_6[T_a], \quad \text{(S44)}$$

$$d_t[T_i] = a_7\Delta^\Gamma[T_i] + \frac{b_7[S](g+[T_a]^2)}{(e+d[T_a]^2)(e+h[W_l])} - c_7[T_i]. \quad \text{(S45)}$$

The activator ($[T_a]$) promotes tentacle initiation, whereas the inhibitor ($[T_i]$) limits its spread, thereby generating a periodic array of tentacle primordia. The system thus provides a minimal description of the tentacle patterning process while remaining functionally decoupled from the Wnt–Dkk axis formation module.

**Extended model for foot formation**

Analogously, the model was further extended to include a subsystem describing foot formation, formulated as a separate activator–inhibitor pair that interacts with the source density ($[S]$) but not with the Wnt–Dkk core system. The foot subsystem represents the basal organiser region and is used to reproduce realistic *Hydra* morphology and regeneration behaviour. The activator is enhanced under conditions of low $[S]$, consistent with the basal identity being promoted in regions distant from the head organiser [13–15].

$$d_t[F_a] = a_8\Delta^\Gamma[F_a] + \frac{b_6(h+[F_a]^2)}{[F_i][S]} - c_8[F_a], \quad \text{(S46)}$$

$$d_t[F_i] = c_9\Delta^\Gamma[F_i] + \frac{b_7(i+[F_a]^2)}{[S]} - d_9[F_i]. \quad \text{(S47)}$$

The foot subsystem is purely phenomenological and does not feed back into the body-axis patterning. It stabilises the aboral pole and contributes to the overall morphological realism of the simulated *Hydra* shape. All parameter values and simulation settings are listed below.

**Numerical implementation pseudo 3D-model**

The mathematical model was simulated using the finite element library Gascoigne [16], based on approximation of the fourth order PDEs in a mixed formulation. We applied linear finite elements for spatial discretisation, and a semi-implicit Euler scheme for time discretisation. For further details of the computation scheme, see [3, 4].

##### Numerical vs. real time and space units

Above mentioned numerical time and space scales can be roughly related to real scales in the following way: The average size of an adult *Hydra* polyp is approximately $X_{real} = 20\,mm$, which corresponds to the length of $X_{num} = 6$ numerical units, leading to the relation $X_{real} \approx \frac{10}{3} X_{num}\,mm$. With respect to time, only a coarse estimation is possible. During simulations, head regeneration is finished for $T_{num} > 0.002$. In experiments, head regeneration requires approx. 48 hours (e.g., [12,17]), leading to $T_{real} \approx T_{num}/(4.2 \times 10^{-5})\,hrs$.

##### Parameters and initial conditions

The mathematical model was simulated using the finite element library Gascoigne [16], which approximates the fourth-order PDEs in a mixed formulation. Linear finite elements were used for spatial discretisation, while a semi-implicit Euler scheme was employed for time discretisation. For further details on the computation scheme, see [3, 4].

Numerical vs. Real Time and Space Units The numerical time and space scales can be roughly related to real-world scales as follows: The average size of an adult Hydra polyp is approximately $X_{\text{real}} = 20$ mm, corresponding to $X_{\text{num}} = 6$ numerical units. This leads to the relation $X_{\text{real}} \approx \frac{10}{3} X_{\text{num}}$, mm. Regarding time, head regeneration in the simulations is completed for $T_{\text{num}} > 0.002$, while experimental data indicate that head regeneration takes approximately 48 hours (e.g., [12,17]), yielding the relationship $T_{\text{real}} \approx \frac{T_{\text{num}}}{4.2\times10^{-5}}$, hrs.

Parameters and Initial Conditions for the Pseudo-3D Model For simulations of the unperturbed system, we used the following parameters for the MI model:

$b_1 = 15 \times 10^{-3}$, $c_1 = 3 \times 10^{-3}$ $a_2 = 18 \times 10^{-7}$, $b_2 = 3 \times 10^{-3}$, $c_2 = 4 \times 10^{-3}$ $a_3 = 22 \times 10^{-3}$, $b_3 = 3 \times 10^{-3}$, $c_3 = 4 \times 10^{-3}$ $a_4 = 1 \times 10^{-7}$, $b_4 = 1 \times 10^{-2}$, $c_4 = 1 \times 10^{-2}$ $a_5 = 11 \times 10^{-6}$, $b_5 = 3 \times 10^{-4}$, $c_5 = 3 \times 10^{-4}$ $d = 0.1$, $e = 1.0$, $f = 0.3$ Additionally, for numerical stabilisation, a small diffusion of the first variable $a_1 = 18 \times 10^{-5}$ was introduced. Model analysis indicated that this did not affect the system dynamics but facilitated numerical approximation of solutions.

For the tentacle and foot pattern formation system, we adapted the following parameters based on Ref. [13, 14, 18]:

$a_6 = 7.5\times10^{-5}$, $b_6 = 2.0\times10^{-2}$, $c_6 = 2.0\times10^{-2}$ $a_7 = 3.0\times10^{-3}$, $b_7 = 3.0\times10^{-2}$, $c_7 = 3.0\times10^{-2}$ $a_8 = 7.2 \times 10^{-4}$, $b_8 = 2.0 \times 10^{-3}$, $c_8 = 3.0 \times 10^{-3}$ $a_9 = 4.4 \times 10^{-2}$, $b_9 = 2.0 \times 10^{-3}$, $c_9 = 3.0 \times 10^{-3}$ $g = 0.005$, $h = 0.01$, $i = 0.0001$ The simulation results were robust with respect to changes in both the qualitative model formulations and parameter variations. Further results, including additional simulations and robustness/sensitivity analysis, are provided in the Supporting Information.

Initial Conditions and Geometry for Hydra Tissue Simulations For the simulations, the geometry of the Hydra tissue was approximated by parametrising initial conditions for $X_1$, $X_2$, and $X_3$ over a closed 2D unit sphere $S^2$ embedded in 3D space, with the initial condition $X_1(t = 0) \equiv X_2(t = 0) \equiv 0$ and $X_3(t = 0) = 4 \cdot s_3$, leading to a stretch in the $s_3$ direction (where $s_1, s_2, s_3$ are Eulerian coordinates of the $S^2$ surface).

For the biological molecules, we used a stochastic initial distribution based on the standard random generator provided by C++. The initial concentration of [S] was modelled using a gradient defined by $[S](t = 0) = 4.0 \cdot (\exp(s_3)/\exp(1))$. For simulations of grafting and head regeneration, $[S](t = 0) = 2.0 \cdot (\exp(s_3)/\exp(1))$ was used, as areas with maximal [S] values were removed during head regeneration. In grafting experiments, initial spots of increased [S] values (mimicking grafted heads) were introduced using circles of radius $r = 0.33$ mm, where $[S] = 4.0$. For simulations of ALP treatment, the initial conditions for the source density were modified to $[S](t = 0) = 2.0 + 4.0 \cdot (\exp(s_3)/\exp(1))$.

To simulate the complete removal of *HyDkk1/2/4-A* or *HyDkk1/2/4-A* expression at a specific time point (after the head pattern had been established), the parameters $b_2$ and $b_4$ were set to zero. HyDkk1/2/4-A knockdown was modeled by increasing $c_2$ from $4 \times 10^{-3}$ to $6 \times 10^{-3}$.

For Hydra aggregates, the initial geometry was a sphere, and all molecules, including [S], were given random initial conditions.

##### Further supporting simulations of the pseudo 3D model

This section contains various supportint simulations of the psuedo 3D model (and derived 1 D simulation profiles) amongst others in comparison to further experimental data.

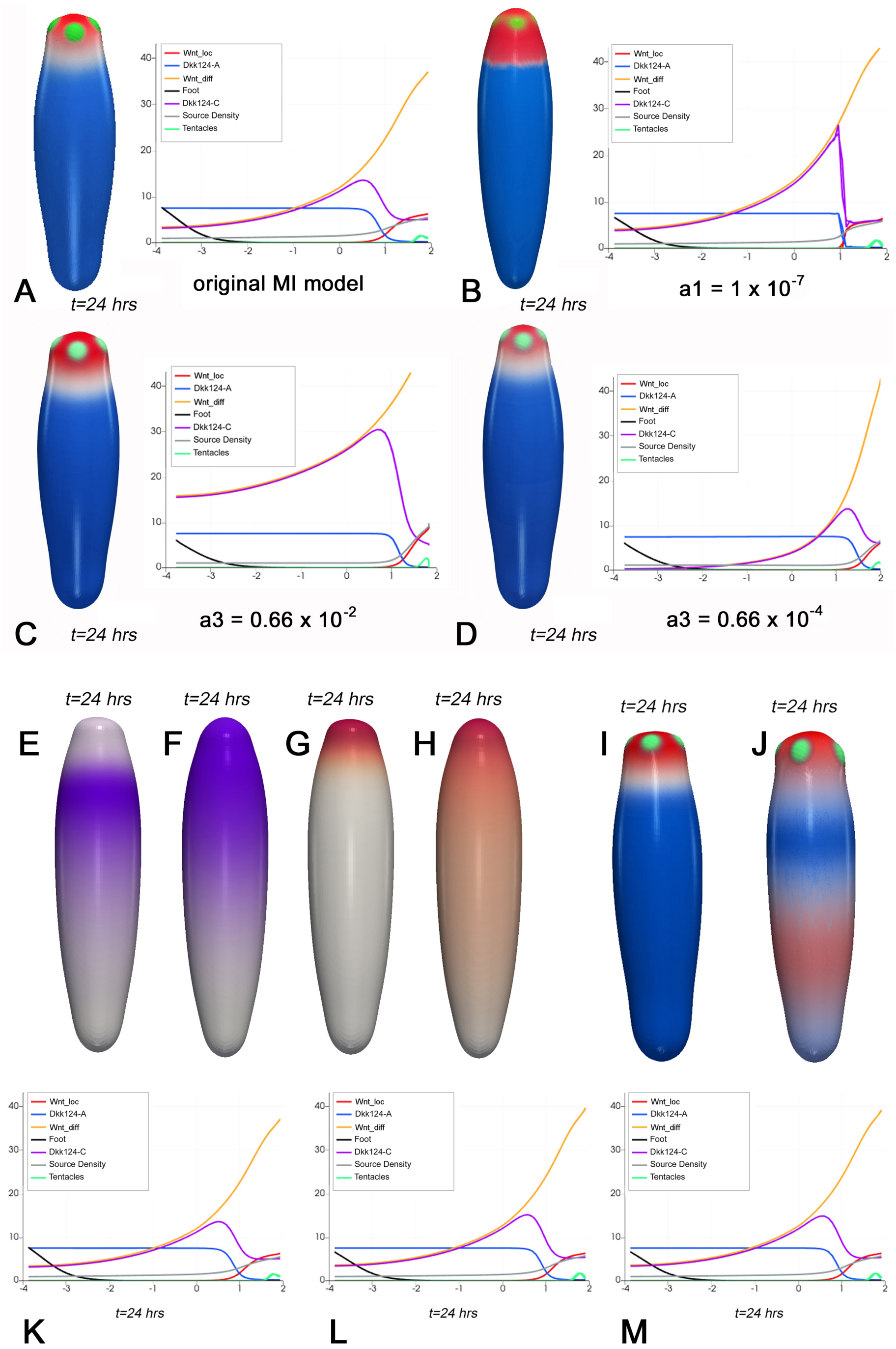


**Fig. S 7.** (A-D): 3D stable patterns and extracted 1D concentration profiles for models with different diffusion rates of the $[W_l]$ complex (B) or of $[W_d]$ (C–D) versus the results for the unperturbed system (A). (E-H): Simulated (rescaled) distribution of Dkk1/2/4-C (E–F) and $[W_l]$ (G–H) expression for the undisturbed system (E,G) vs. a system in which no $[W_l]$-based inhibition of Dkk1/2/4-C has been considered (F,H). In particular, in the latter, expression patterns resemble the classical activator-inhibitor model. (I-J): Relative distribution of simulated Dkk1/2/4-A (blue), $[W_l]$ (red), and tentacle (green) expression for the undisturbed system (I) vs. a system without constant *Dkk1/2/4-A*-expression but an [S]-induced expression instead (J). (K-M): Concentration profiles of the unperturbed system (K) versus the system with equal Dkk1/2/4-A and Dkk1/2/4-C diffusion rates (L–M). (L): $a_2 = a_4 = 1 \times 10^{-6}$: (M): $a_2 = a_4 = 1 \times 10^{-7}$.

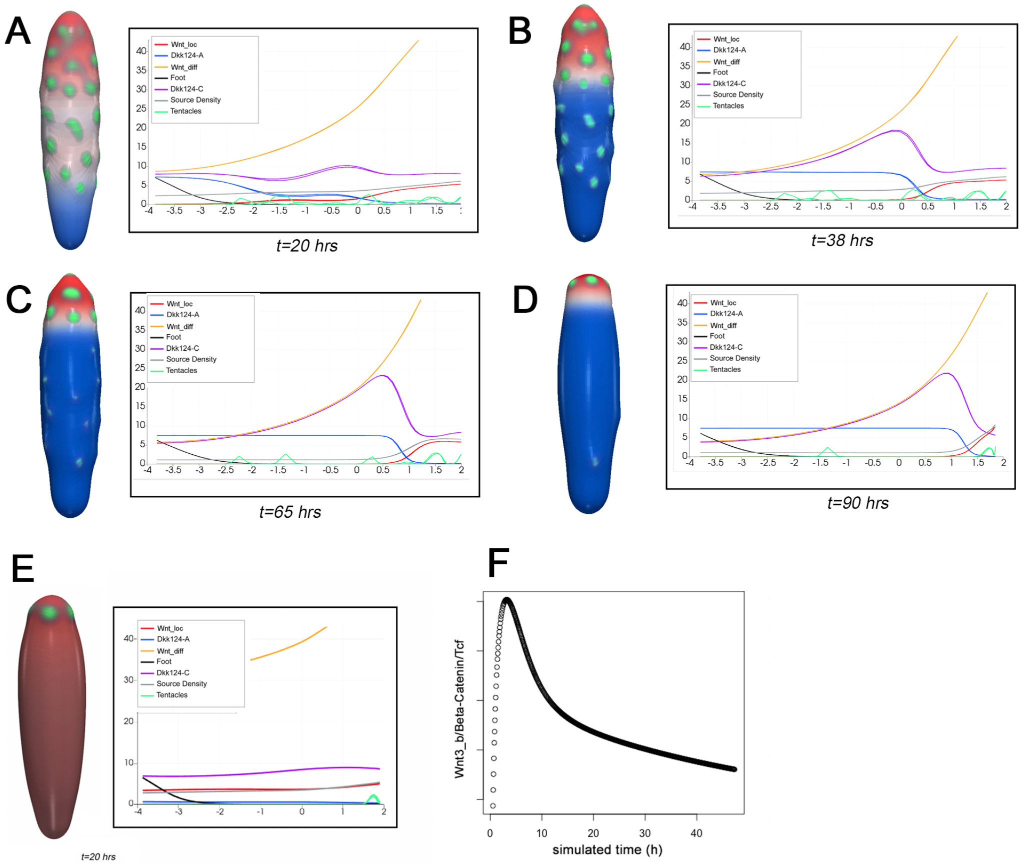


**Fig. S 8.** (A–D): 3D results and extracted 1D concentration profiles for different time points after simulated ALP treatment. Red colour represents the $[W_l]$ complex, blue colour Dkk1/2/4-A and green colour tentacles. (E): Snapshot of a simulation without the Dkk1/2/4-A-based inhibition of the $[W_l]$ complex. Observed patterns (e.g., $Wnt3_{diff}$ gradients) result from [S]-initial conditions only, and vanish over time. I.e., the mutual negative feedback loop is required for local self-activation. (F): Simulated temporal development of relative $\beta$-Catenin expression strength after head removal.

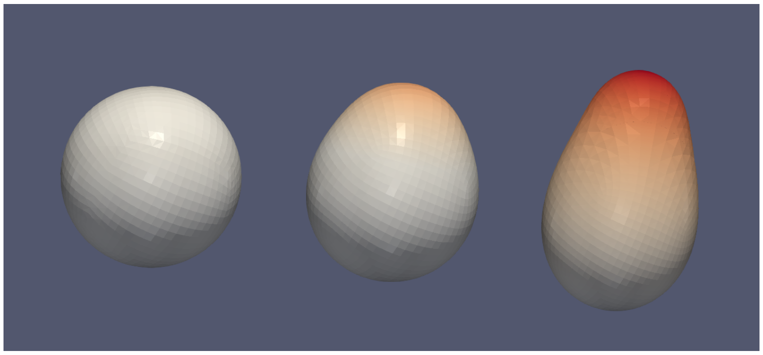

**Fig. S 9.** Different stages of the 3D simulation of the *Hydra* aggregate system without foot and tentacle modules. The simulation corresponds to Fig. 2D in the main text but excludes both auxiliary pattern-formation subsystems. The resulting *de novo* symmetry breaking and Wnt–Dkk distributions remain unchanged (apart from the absence of foot and tentacle structures), confirming that the foot and tentacle modules are purely morphological features and do not affect axis patterning.

To verify that the additional modules for foot and tentacle formation do not influence axis patterning, we repeated the 3D simulation of the *Hydra* aggregate system shown in Fig. 2D of the main text with both modules removed. The resulting *de novo* symmetry breaking and steady-state axis pattern were indistinguishable from the full model. This confirms that the foot and tentacle systems are purely morphological extensions and have no feedback on the Wnt–Dkk mechanism responsible for axis formation (cf., Fig. S 9).

#### Experimental and conceptual background of model components

The following section provides the biological and experimental background supporting the structure of the model equations presented in the main text. Each equation term corresponds to experimentally observed regulatory interactions or theoretical concepts that have been established in previous studies on *Hydra* axis formation and patterning.

*Cell-local Wnt dynamics (Eq. 1):* The variable $[W_l]$ represents intracellular canonical Wnt activity, summarizing the combined effects of $\beta$-catenin/Tcf signalling and expression of one or more canonical *HyWnt* genes. The production of $[W_l]$ is promoted by the source density ($[S]$), which defines the tissue's positional competence. This formulation reflects the experimental observation that tissues with higher head-forming capacity are more prone to activate Wnt/$\beta$-catenin signalling [13, 19–21]. The two Dkk proteins (HyDkk1/2/4-A and HyDkk1/2/4-C) act as inhibitors of this activity [22, 23], consistent with their mutually repressive interaction with canonical Wnt signalling. The remaining denominator term introduces a natural saturation of production, preventing unbounded activation, while the final term in Eq. (1) accounts for degradation or turnover of Wnt-related activity. Together, these terms reproduce the experimentally observed localization of strong Wnt activity at the apical tip and its suppression in the body column [11, 17, 21, 24–26].

*Dkk1/2/4-A dynamics (Eq. 2): HyDkk1/2/4-A* ($[A]$) acts as a short-range inhibitor of canonical Wnt signalling. Its expression is repressed by local Wnt activity ($[W_l]$), consistent with alsterpaullone (ALP) treatment experiments showing that *HyDkk1/2/4-A* expression is negatively regulated by Wnt/$\beta$-catenin activity and that its removal leads to ectopic *HyWnt3*-expressing clusters along the body column [22]. The inhibitory influence of $[A]$ on Wnt signalling has been confirmed both in *Hydra* and in *Xenopus* assays [22], and its knockdown combined with ALP treatment induces ectopic axis formation [27]. Eq. (2) therefore includes basal production repressed by $[W_l]$, small diffusion to mediate short-range inhibition, and a degradation term for turnover. Together, these interactions capture the experimentally observed mutual inhibition between *HyDkk1/2/4-A* and canonical Wnt signalling.

*Diffusible Wnt dynamics (Eq. 3):* The diffusible Wnt variable, $[W_d]$, represents secreted Wnt ligands that spread over several cell diameters and mediate longer-range feedback. Its production depends on both $[S]$ and $[W_l]$, expressed as the multiplicative term $[S] \times [W_l]$. This coupling reflects the biological requirement that Wnt transcription and secretion occur only where tissue is both competent (high $[S]$) and actively signalling via $\beta$-catenin/Tcf (high $[W_l]$). $\beta$-Catenin/Tcf alone regulates thousands of genes [28] and is therefore not sufficient to specify Wnt gene expression or secretion. Additional slower regulatory layers—such as chromatin accessibility, stable cellular states, or other long-term determinants of gene expression—likely define whether a region is permissive for head-related gene expression. The $[S]$ field can thus be interpreted as a coarse-grained representation of such permissive conditions, while $[W_l]$ provides the immediate transcriptional drive. Only where both factors coincide can secreted Wnts be produced, ensuring robust and spatially confined activation of the head organizer. The diffusion of $[W_d]$ along the epithelial surface represents the lateral spread of secreted molecules, enabling intercellular coupling across the tissue. While studies in other organisms indeed suggest such diffusion [29–31], direct evidence in *Hydra* is still lacking. However, simulation results are robust over a broad range of diffusion constants (see Supplementary Fig. S6A–D), and the qualitative patterning behaviour does not depend critically on these values. Finally, the last term in Eq. (3) accounts for degradation or clearance of secreted Wnt molecules.

*Dkk1/2/4-C dynamics (Eq. 4): HyDkk1/2/4-C* ($[C]$) represents a second inhibitory component with distinct regulation from $[A]$. Its production is positively influenced by diffusible Wnt ($[W_d]$) and negatively regulated by local Wnt activity ($[W_l]$), consistent with co-expression and LiCl treatment experiments [23]. This dual control ensures that $[C]$ is activated by long-range Wnt signals but suppressed within the head organizer itself, thereby restricting the spatial domain of Wnt activity. The broader spatial expression domain of *HyDkk1/2/4-C* compared to *HyWnt3*, as well as its characteristic ring-like pattern around the head region, suggests that activation and inhibition act on different spatial scales [18, 23]. This supports the inclusion of long-range activation by secreted Wnt ligands in Eq. (4). Removal of *HyDkk1/2/4-C*-expressing cells leads to extension of the *HyWnt3* domain into the head region [23], confirming its inhibitory role. Eq. (4) also includes small diffusion and degradation terms, reflecting the limited range and turnover of this inhibitor.

*Source density dynamics (Eq. 5):* The source density ($[S]$) captures a slowly varying, long-term field that encodes positional information along the oral–aboral axis of *Hydra*. This variable corresponds to the experimentally observed "head-forming capacity" or positional competence, which persists for several days after grafting or regeneration and depends on canonical Wnt/$\beta$-catenin activity [13, 19–21, 24]. Eq. (5) combines diffusion, Wnt-dependent production, and slow decay. Diffusion reflects the gradual redistribution of competence across the tissue, the production term represents induction of long-term competence by sustained Wnt/$\beta$-catenin activity [32], and the decay term models its relaxation back to baseline levels. Conceptually, $[S]$ can be interpreted as an emergent property arising from slow molecular, cellular, or possibly epigenetic mechanisms that determine which regions are permissive for head-related gene expression and organizer formation. This abstraction allows the model to capture the experimentally observed stability and spatial organization of the head-forming capacity without assuming a specific molecular identity.

In summary, each equation term is directly linked to experimental observations or well-established theoretical principles. The resulting system captures the interplay between short-term signalling (Wnt/$\beta$-catenin, Dkk feedback) and long-term positional memory (source density), providing a unified description of axis patterning in *Hydra*.

29. Mii Y, Nakazato K, Pack CG, Ikeda T, Sako Y, Mochizuki A, et al. Quantitative Analyses Reveal Extracellular Dynamics of Wnt Ligands in Xenopus Embryos;10:e55108. doi:10.7554/eLife.55108.

30. Pani AM, Goldstein B. Direct Visualization of a Native Wnt in Vivo Reveals That a Long-Range Wnt Gradient Forms by Extracellular Dispersal;7:e38325. doi:10.7554/eLife.38325.

31. Recouvreux P, Pai P, Torro R, Ludányi M, Mélénec P, Boughzala M, et al.. Establishment of Wnt Ligand-Receptor Organization and Cell Polarity in the C. Elegans Embryo;. Available from: `https://www.biorxiv.org/content/10.1101/2023.01.17.524363v1`.

32. Tursch A, Bartsch N, Mercker M, Schlüter J, Lommel M, Özbek AMCA, et al. Injury-induced MAPK activation triggers body axis formation in Hydra by default Wnt signaling. PNAS. 2022;119(35):e2204122119.

### Supporting Information Legends

#### S1 Text. One-dimensional model reduction

Detailed derivation of the nondimensionalised form of the one-dimensional model, including stability analysis of spatially homogeneous steady states and discussion of conditions leading to Turing instability (see Supporting Information, pp. 1–5).

#### S1 Table. Parameter values of the one-dimensional model

Table summarising fitted parameter values $\nu_i$, $\beta_i$, and $\tau_i$ for the reduced one-dimensional model (Table S1, p. 5).

#### S1 Fig. Intersections of $f_1(w)$ and $f_2(w)$

Illustration of steady-state structure for different values of $\beta_1$, showing the number and location of intersections between $f_1(w)$ and $f_2(w)$ (Fig. S1, p. 3).

#### S2 Fig. Comparison of 1D steady-state patterns with 3D data

Steady-state concentration profiles of the one-dimensional model compared with averaged profiles extracted from 3D simulations. Both patterns are scaled using min–max normalisation (Fig. S2, p. 6).

#### S3 Fig. Examples of pattern formation for different initial conditions

Three examples of pattern formation in the one-dimensional model for different initial perturbations of the homogeneous steady state (Fig. S3, p. 6).

#### S4 Fig. Bifurcation and eigenvalue analysis

Eigenvalue structure of the linearised operator at the critical diffusion value, unstable modes as functions of diffusion, and comparison between asymptotic and numerically obtained stationary solutions (Fig. S4, p. 10).

#### S5 Fig. Pattern of modified model with feedback in degradation terms

Example pattern obtained from numerically integrating the model variation defined in Eqs. (S34)–(S38), which introduces feedback via modified degradation terms (Fig. S5, p. 10).

#### S6 Fig. Pattern of model variant with nonlinear reaction terms

Numerical pattern obtained from the model using higher-order reaction terms in Eqs. (S39)–(S43), demonstrating robustness of the patterning mechanism (Fig. S6, p. 11).

#### S7 Fig. Parameter sensitivity analysis

Comparison of stable 3D patterns and extracted 1D profiles for different diffusion rates and inhibition conditions, including simulations with altered Wnt–Dkk interactions (Fig. S7, p. 15).

#### S8 Fig. ALP-treatment simulations

Simulation results showing temporal pattern development following ALP treatment, comparison with a system lacking Dkk1/2/4-A inhibition, and simulated $\beta$-catenin recovery dynamics (Fig. S8, p. 16).

#### S9 Fig. Symmetry breaking in aggregate simulations without auxiliary modules

Stages of axis formation in Hydra aggregates in the absence of tentacle and foot subsystems, demonstrating that these modules do not influence Wnt–Dkk axis patterning (Fig. S9, p. 17).
